# Somatostatin venom analogs evolved by fish-hunting cone snails: From prey capture behavior to identifying drug leads

**DOI:** 10.1101/2021.10.26.465842

**Authors:** Iris Bea. L. Ramiro, Walden E. Bjørn-Yoshimoto, Julita S. Imperial, Joanna Gajewiak, Maren Watkins, Dylan Taylor, William Resager, Beatrix Ueberheide, Hans Bräuner-Osborne, Frank G. Whitby, Christopher P. Hill, Laurent F. Martin, Amol Patwardhan, Gisela P. Concepcion, Baldomero M. Olivera, Helena Safavi-Hemami

## Abstract

Somatostatin (SS) is a peptide hormone with diverse physiological roles. By investigating a deep-water clade of fish-hunting cone snails, we show that predator-prey evolution has generated a diverse set of SS analogs, each optimized to elicit specific systemic physiological effects in prey. The increased metabolic stability, distinct SS receptor activation profiles, and chemical diversity of the venom analogs make them suitable leads for therapeutic application, including pain, cancer and endocrine disorders. Our findings not only establish the existence of SS-like peptides in animal venoms, but also serve as a model for the synergy gained from combining molecular phylogenetics and behavioral observations to optimize the discovery of natural products with biomedical potential.

**One Sentence Summary:** Somatostatin drug design by fish-hunting cone snails

## Main Text

Cone snails comprise a large family of venomous marine predators (family *Conidae*). The complete lack of offensive mechanical weaponry in this hyperdiverse molluscan lineage has been compensated for by the evolution of highly sophisticated pharmacological strategies for prey capture (*1*). Because of their stability, chemical diversity, and target selectivity, cone snail toxins (conotoxins) have been developed as biomedical tools (*2–4*), drug leads, and drugs (*5-8*), and have elucidated previously unknown signaling pathways in health and disease (*2, 9-11*).

The ~1000 living species of cone snails can be broadly divided into those that hunt fish, gastropod mollusks, or marine worms (primarily polychaetes) (*1*). Because many of the molecular targets expressed in fish also play analogous important physiological roles in humans, fish-hunting *Conus* species have been the focus of many cone snail venom research programs. Eight lineages of fish hunters exist, comprising approximately 150 extant species (*12*). A phylogenetic reconstruction of representatives of these lineages is shown in Fig. 1A. Within these, diverse predation strategies have evolved for capturing fish prey.

**Fig. 1.**
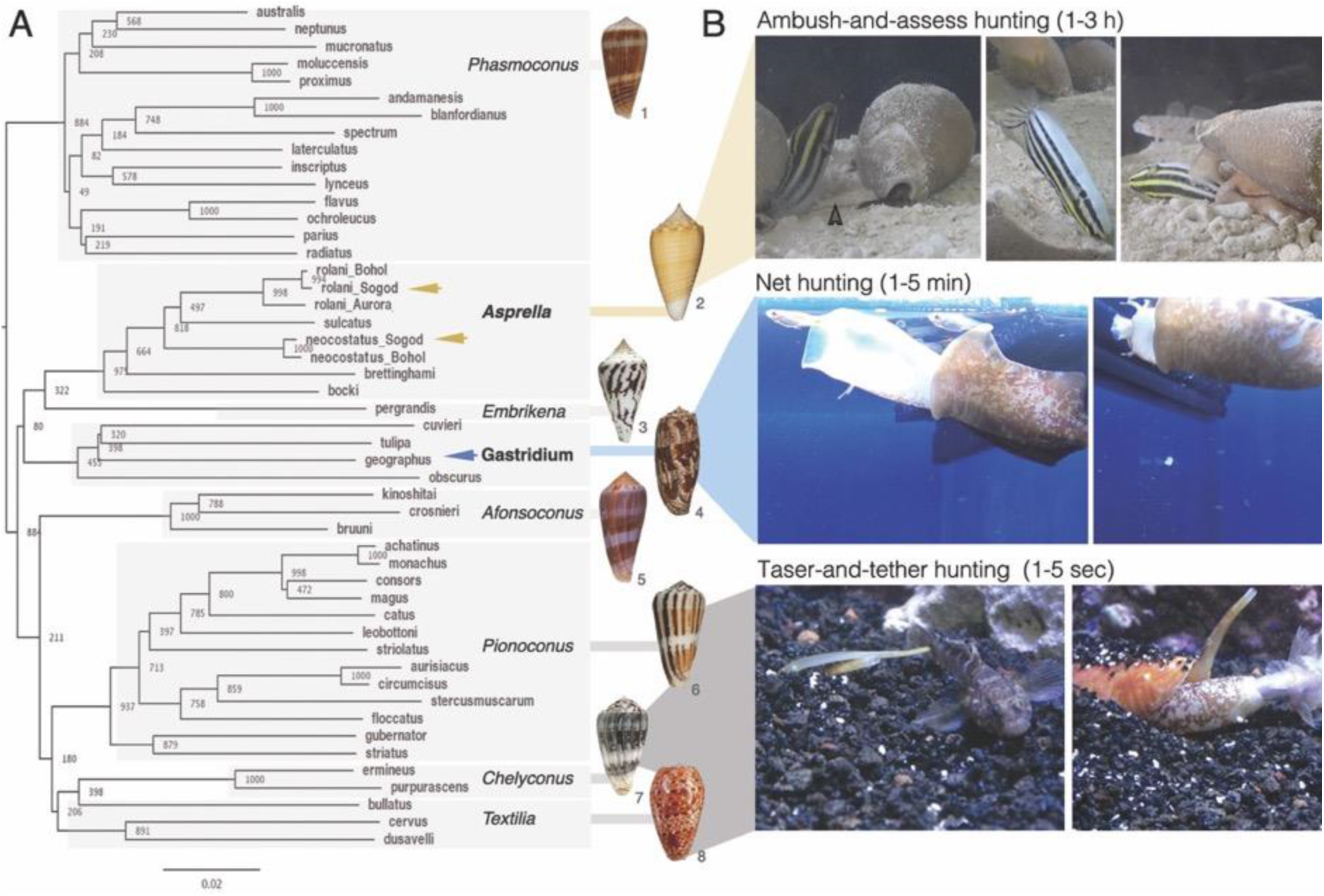
Evolution of distinct predation strategies in different clades of fish-hunting cone snails. **A.** Phylogenetic tree of the eight recognized clades of fish hunters. Bootstrap values and shell images of neotypes are shown (1: *Conus radiatus*, 2: *Conus sulcatus*, 3: *Conus pergrandis*, 4: *Conus geographus*, 5: *Conus kinoshitai*, 6: *Conus magus*, 7: *Conus ermineus*, 8: *Conus bullatus*). **B.** Still images of three distinct predation strategies filmed for three different species of fish hunters (from top to bottom: *C. neocostatus* (*Asprella* clade, arrow marks proboscis loaded with radula tooth), *C. geographus* (*Gastridium* clade), *C. bullatus*, (*Textilia* clade)). Images were extracted from supporting movie files (Movies S1-3).

The most widespread and best studied fish-hunting strategy is called taser-and-tether (*13*). First described in 1956, it is characterized by a rapid chemical electrocution of prey that is facilitated by the action of diverse ion channel modulators (Movie S1, Fig. 1B) (*13*). The underlying molecular mechanisms of taser-and-tether toxins have been well characterized and have provided a rich set of pharmacological tools and drug leads, including an approved therapeutic for the treatment of chronic pain (*5*).

A second predation strategy, net hunting, has been observed in only two species, *Conus geographus* and *Conus tulipa* from the Gastridium clade. Behavioral observations dating back to the early 1970s showed that the net hunting strategy is characterized by a pre-capture phase during which venom components released into the water cause fish to become both sensory deprived and disoriented, thereby facilitating prey capture (*1, 14*). The mixture of compounds that elicit this response were called the nirvana cabal because they make fish appear as if under the influence of narcotic drugs (*1*). We provide a video recording showing that the net-hunting strategy enables simultaneous capture of multiple fish (Movie S2, Fig. 1). Nirvana-cabal toxins have proven biomedical applications, including a diagnostic tool for an autoimmune disorder (*15*) and a family of insulins that have inspired the design of fast-acting drug leads for diabetes (*16, 17*). *Conus geographus* venom has also provided one of the most widely-used pharmacological tools in molecular neuroscience for studying synaptic transmission, the calcium channel blocker ω-conotoxin GVIA (*5*).

In this article, we document the discovery that a largely unexplored, deep-water lineage of fish-hunting cone snails of the *Asprella* clade uses a strikingly-different predation strategy that we call “ambush-and-assess”. In contrast to the rapid and efficient capture previously described, direct observation of the *Asprella* envenomation behavior demonstrates that it takes 1-3 hours from the first strike to engulfment of prey. This unexpected observation led us to investigate venom components that potentially facilitate the novel, idiosyncratic behavior observed. By combining the behavioral observations of prey capture by the cone snail predator with behavioral changes elicited by the action of individual venom components in mice, a novel class of analogs of the vertebrate hormone somatostatin was discovered.

## Results and Discussion

### Deep-water cone snail species of the *Asprella* clade: observations of fish-prey envenomation

Phylogenetic reconstruction using the *mitochondrial cytochrome* c oxidase subunit 1 (*COI*) marker gene highlights the existence of eight lineages of fish hunters in the genus *Conus* (Fig. 1A) (*12, 18*). Of these, the *Asprella* clade is the least well-known (*1*). Unlike the majority of cone snail species that inhabit shallow waters, *Asprella* species live well offshore, typically at depths of 60 – 250 m (*19*). This has hampered behavioral observation and venom analysis. To overcome this limitation, we established contacts with fishermen experienced in collecting marine gastropods from deep waters off the town of Sogod on Cebu Island in the Philippines. After a series of field expeditions, we collected enough specimens of two species of *Asprella, Conus rolani* and *Conus neocostatus*, for behavioral recordings and bioactivity-guided toxin discovery.

Three captive individuals of *C. neocostatus* were successfully documented catching different species of fish (Movies S3-5, see Table S1 for timeline of events). In all three predation episodes, *C. neocostatus* struck its prey with a rapid venom sting followed by immediate withdrawal (< 1 s). Following envenomation, the fish initially appeared normal and continued to swim for 5-15 minutes. The snail waited during this time and did not react to any other fish that may have approached. In two of the three cases, the cone snail approached the fish again and carried out a second strike (Movie S4 and S5). In all three cases, once the fish was deemed to be dead or completely incapacitated, the snail approached the fish from behind and consumed it whole. The timeline from first strike to the quiescent fish being engulfed ranged from 1 - 3.5 h. Thus, in striking contrast to prey capture by fish-hunting cone snails of other phylogenetic lineages, prey capture by *Asprella* is extremely slow.

We hypothesize that this seemingly inefficient predation behavior enables capture of aggressive fish that need to be approached from a distance and cannot be tethered without risk of retaliation. In line with this, we have documented fish that aggressively attack cone snails from other lineages, as they are extending their proboscis in preparation to envenomate their prey (Movie S6). Killing of potentially dangerous prey has been proposed as one of the most significant factors of natural selection on predatory organisms (*20*). Notably, predation by *Asprella* snails bears resemblance to the strike-and-release strategy used by rattlesnakes and other viperids of the family *Viperidae*, and by some of the deadliest members of the *Elapidae* family, including cobras (*20*). These snakes ambush rodent prey with an extremely rapid envenomating strike followed by a quick release of prey (*21*). The envenomated rodent then wanders off before ultimately succumbing to the deadly sting. Envenomation without capture provides the advantage of minimal contact with dangerous prey but adds the risk of potentially losing the prey and the challenge of relocating the envenomated animal. After envenomation, even when multiple fish were available, *C. neocostatus* tracks the individual that it struck, possibly indicating that the snail can detect chemosensory cues released by the envenomated prey as has been reported for rattlesnakes (*22*).

Anecdotal evidence suggests that ambush-and-assess hunting has also evolved in other cone snail lineages. *Conus flavus*, a fish-hunting species of the *Phasmoconus* clade, reportedly envenomates and captures fish without tethering (*1*). However, in contrast to *Asprella*, envenomation by *C. flavus* leads to relatively faster immobilization. The extremely slow onset of action observed here suggests that the venom of *Asprella* snails may contain toxins that target G protein-coupled receptors (GPCRs) expressed in the somatosensory or neuroendocrine systems of prey. To specifically identify these toxins, we performed bioactivity-guided assays with an emphasis on compounds that elicited behavioral changes reminiscent of the slow-onset state of hypoactivity observed in fish after *Asprella* envenomation.

### *Asprella* venom elicits a slow onset hypoactivity in mice; identification of a novel bioactive peptide

Venom was extracted from venom glands of *Conus rolani*, the *Asprella* species that is collected in the deepest water habitats (200-1250 m) (*19*). Unlike *C. neocostatus, C. rolani* does not survive in captivity but can be abundantly collected, making the accumulation of larger quantities of venom for activity testing possible. Because of their well-established use in toxinology and their closer relatedness to humans, mice were used to assay venom activity instead of fish. Activity testing of venom fractionated by reversed-phase chromatography revealed the presence of a fraction that elicited a phenotypic response upon intracranial (IC) injection in mice that was very similar to what was observed in fish: slow-onset loss of balance and a state of hypoactivity and sedation that lasted for > 3 h (fraction #16, Fig. 2A, Table S2). Notably, this fraction contained several toxins including an insulin-like peptide and a member of the conantokin toxin family (data not shown). However, these two components were not further pursued because of their similarity to known cone snail toxins (*23, 24*). Instead, we focused our discovery efforts on a third compound with yet unknown chemistry and biology.

**Fig. 2.**
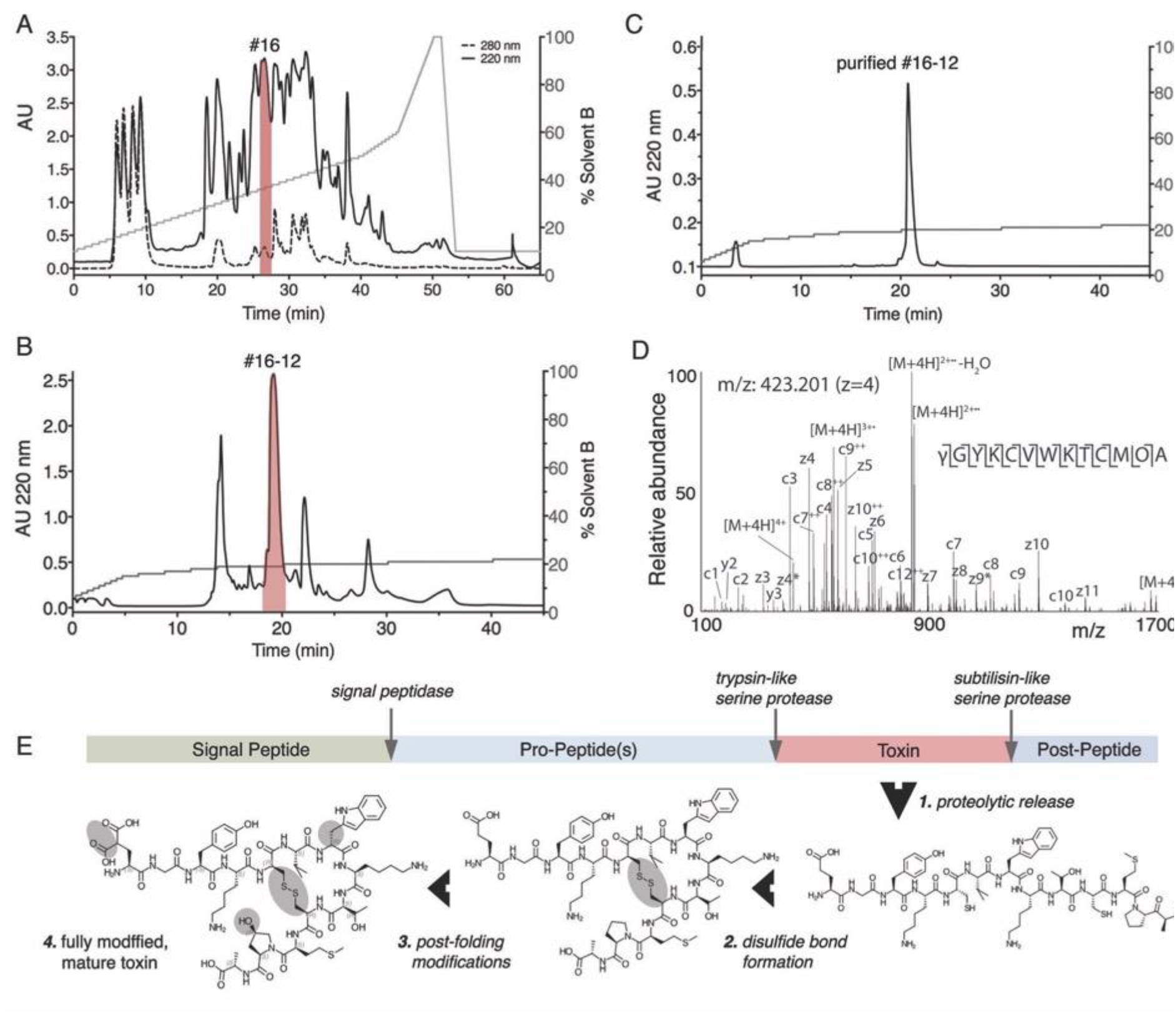
Identification of Consomatin Ro1. **A.** Reverse-phase chromatography of crude venom of the *Asprella* snail *C. rolani* and **B.** subfractionation and **C.** purification of Consomatin Ro1 from subfraction 16#12 (shown in red). **D.** Electron Transfer Dissociation (ETD) MS/MS spectrum of the quadruple charged ion of the venom peptide after reduction and alkylation with 2-methylaziridine acquired on the Orbitrap Lumos Tribrid with 30,000 resolution (@ 400 *m/z*) and 3 micro scans. *N*-terminal fragment ions (c-type ions) are indicated by ⌉ and C-terminal fragment ions (z,y-type ions) are indicated by ⌊. Doubly charged ions are indicated with ++ and z ions resulting from cleavage at cysteine and loss of the cysteine side chain are indicated with *. [M + 4H]^2+••^ and [M + 4H]^1+•••^ indicates charge reduced species. Due to space limitations, not all different charge states of already labeled peptide bond cleavages are indicated in the figure. The mass accuracy for all fragment ions is better than 10 ppm. γ= γ-carboxyglutamate; O= hydroxyproline; w=D-tryptophan. **E.** Organization of the prepropeptide of Consomatin Ro1 identified by transcriptome sequencing depicting post-translational intermediates and processing events.

The relevant venom fraction was further fractionated, and the bioactivity of individual components assessed by injecting them in mice. Subfraction #16-12 (Fig. 2B-C) most closely recapitulated the initial effect observed in fraction 16 (Table S2). The active component of subfraction #16-12 was characterized using a combination of Edman degradation, mass spectrometry (MS) and transcriptome sequencing. As determined by MS, the compound that elicited the bioactivity was a small peptide with a monoisotopic mass of 1573.65 (M+1H)^1+^ (Fig S1). This peptide is hereafter referred to as Consomatin Ro1.

### Biochemical characterization and bioactivity of Consomatin Ro1

Tandem MS sequencing of purified Consomatin Ro1 and database searches of MS spectra against the venom gland transcriptome of *C. rolani* revealed the peptide’s amino acid sequence and post-translational modifications (Fig. 2D). In addition to a disulfide bond between Cys5 and Cys10, the 13-residue peptide contains an N-terminal γ-carboxylated Glu (abbreviated as γ) and a hydroxylated Pro (abbreviated as O) in position 12 (Fig. 2D-E). Furthermore, when sequenced by Edman degradation, Trp7 gave a lower-than-expected signal, indicating that this amino acid was potentially modified to a dextrorotary (D)-Trp. In order to investigate the chirality of Trp7, we synthesized two analogs of Consomatin Ro1 containing either the L-Trp7 or the D-Trp7 enantiomer and compared their reversed-phase elution profile and *in vivo* activity to that of the native venom peptide. Only the D-Trp7 analog co-eluted with the venom peptide (Fig. S2). Additionally, Consomatin Ro1 (D-Trp7) exhibited the same bioactivity as the purified venom peptide: mice behaved normally for the first few minutes after IC injections but became progressively less active, and ultimately motionless. These effects were dose-dependent, with the lowest dose needed to elicit these effects being 0.52 - 0.94 mg/kg body weight (Table S2). At 4 - 5 hours post-injection mice recovered, even from the highest dose tested (5.4 mg/kg).

Transcriptome sequencing of the *C. rolani* venom gland identified the full-length transcript encoding Consomatin Ro1. The translated Consomatin Ro1 precursor sequence belongs to the C-gene superfamily and shares the canonical sequence organization of most cone snail toxins, consisting of an N-terminal signal sequence for translocation into the endoplasmic reticulum and secretory pathway, followed by the propeptide region, the mature toxin, and a short postpeptide sequence (Fig. 2E and Fig. S3). Consomatin Ro1 appears to be released from the prepropeptide by trypsin-and subtilisin-like serine proteases. The biological function of the pre- and post-peptide region of the Consomatin Ro1 precursor remains to be determined but given the high degree of modifications, may include recognition sites for modifying enzymes.

### Consomatin Ro1 is a mimetic of the vertebrate hormone somatostatin

The slow onset of action of Consomatin Ro1 in mice suggested that this peptide may not be a classical ion channel modulator but might instead target a GPCR. Indeed, comparative sequence mining of Consomatin Ro1 against an in-house database of known GPCR ligands revealed the toxin’s similarity to somatostatin (SS), an agonist of the *G protein*-coupled SS receptor family (Fig. 3).

**Fig. 3.**
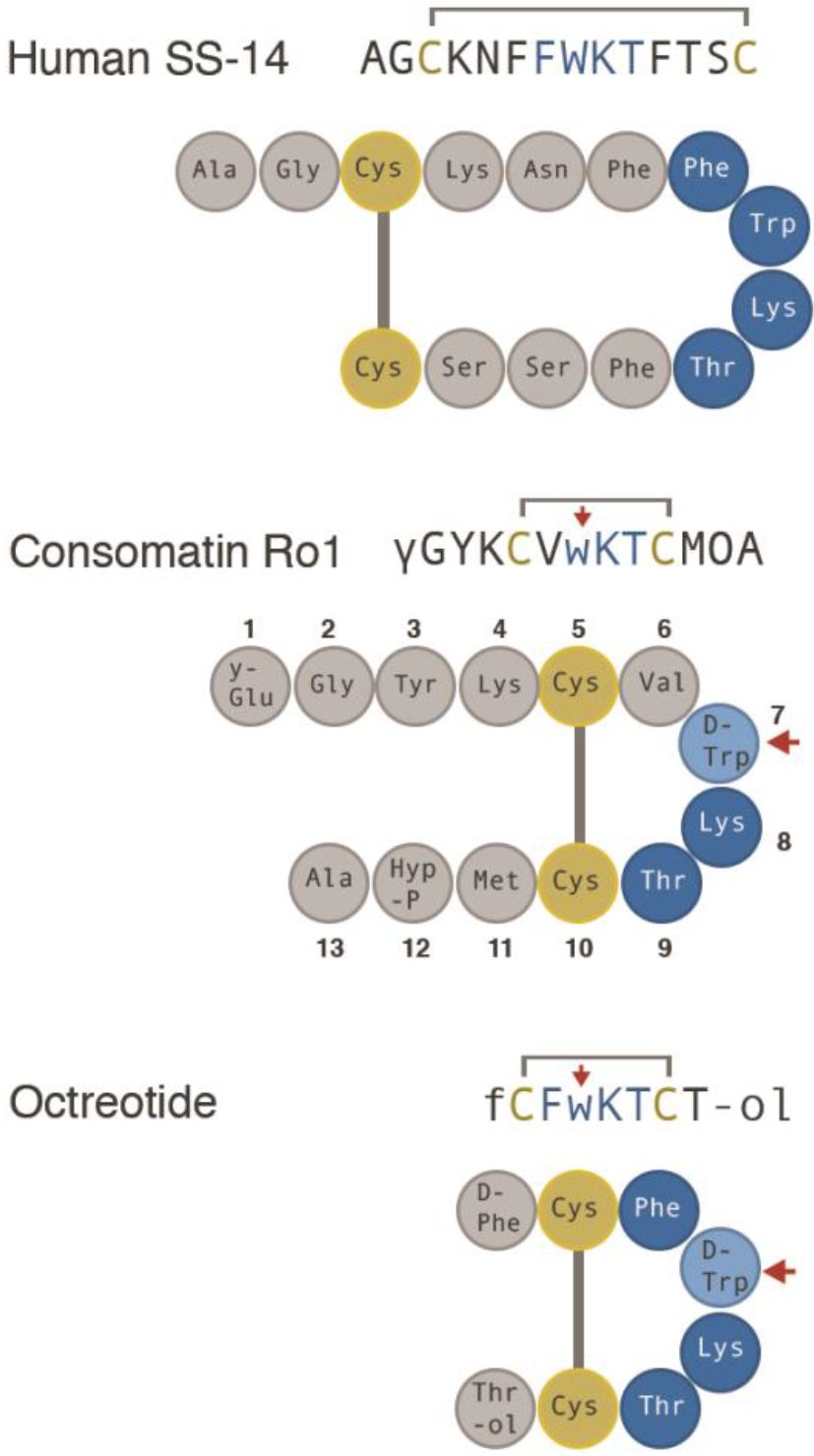
Sequences and schematic representation of human SS, Consomatin Ro1, and the SS drug analog Octreotide. Cysteines and amino acids of the core SS receptor binding motif are shown in yellow and blue, respectively. Modifications: γ=γ-carboxyglutamate; w=D-tryptophan (marked by red arrow); O=hydroxyproline; f=D-phenylalanine; #=amidation; -ol=alcohol.

SS is a highly conserved vertebrate peptide hormone that, in humans, is predominantly secreted in the brain, pancreas, and gastrointestinal tract where it functions as an inhibitor of secretion and cell proliferation (*25*). SS was first discovered as the main inhibitor of growth hormone (GH)-release from the anterior pituitary gland and has since been associated with many additional biological functions, including the inhibition of pancreatic hormone secretion, neuronal signaling, pain, and inflammation (*26*). SS belongs to a larger family of related peptide hormones that includes SS-14, SS-28, and cortistatin. All three peptides bind to and activate the five subtypes of the SS receptor (SST_1-5_ (*27*)). Notably, the venom peptide Consomatin Ro1 shares a disulfide bond and three of the four core residues known to be important for SS receptor activation (Trp7-Lys8-Thr9; numbering according to the sequence of Consomatin Ro1 throughout the manuscript) (*28*) (Fig. 3).

To the best of our knowledge, Consomatin Ro1 represents the first SS-like peptide to be isolated from venom. Additionally, while SS is ubiquitously found in vertebrates, including fish (*29*), and has recently also been reported in invertebrate deuterostomes (*30*), the presence of a peptide with such a high degree of sequence similarity to SS in a snail is surprising since the existence of SS-like peptides in protostomes is still a matter of controversy (*29, 31*).

In addition to SS, Consomatin Ro1 shares some sequence similarity with the peptide hormone urotensin-II (UII). UII activates the urotensin receptor (UT), another class A GPCR, and functions as one of the most potent vasoconstrictors in humans (*32*). However, UT-II lacks several residues important for SS receptor activation, and has low activity at SST_1-5_ (*32*).

### Consomatin Ro1 is an evolutionarily optimized stable analog of SS

Given its important roles in human physiology, SS has been investigated as a drug lead for various diseases, including hypersecretory tumors, diabetes, cancer, pain, and inflammation (*28*). Early attempts to develop SS as a drug proved unsuccessful because of the hormone’s very short *in vivo* half-life and lack of receptor subtype selectivity (*33*). These limitations fueled efforts to design metabolically-stable, subtype-selective SS analogs (*34*). The three main features that are commonly shared between these analogs are a modified N-terminus, a shortened cyclic core, and the incorporation of a D-Trp within the conserved Phe-Trp-Lys-Thr receptor binding motif (Fig. 3). These modifications stabilize a characteristic pharmacophoric β-turn motif, provide increased resistance to proteolytic degradation, and confer subtype selectivity (*34*). For example, the *in vivo* half-life of Octreotide (tradename Sandostatin^®^), an SS drug analog approved for the treatment of GH-producing tumors, is 90 min following intravenous infusion compared to 1-3 min for native human SS (*35, 36*).

Remarkably, the venom peptide Consomatin Ro1 not only displays all of the three drug-like features introduced into minimized therapeutic SS analogs, but both the disulfide bond and the D-Trp are located at the same position (Fig. 3). Consistent with this, the *in vitro* half-life of Consomatin Ro1 when incubated with human plasma at 37 °C is >158 h compared to 5.5 h for native human SS (Fig. S4). While incorporation of D-enantiomers is a common strategy used for enhancing the metabolic stability of compounds in drug design (*37*), in nature, D-amino acid containing peptides and proteins are extremely rare, particularly in metazoans (*38*). To the best of our knowledge, no other natural product has ever been reported to share a D-enantiomer with a synthetic peptide drug.

### The X-ray structure of Consomatin Ro1 closely aligns with that of the SS drug analog Octreotide

In order to compare Consomatin Ro1 to SS and therapeutic SS drug analogs, we determined the X-ray structure of this toxin at 1.95 Å resolution. Crystallographic data and refinement statistics are provided in Table S3. The structure of Consomatin Ro1 is characterized by two β-strands and a β-turn in the cyclic core (residues Phe6-D-Trp7-Lys8-Thr9) between the Cys5-to-Cys10 disulfide linkage (Fig. 4A). The crystal of Consomatin Ro1 contained 6 molecules per asymmetric unit, arranged in an anti-parallel β-sheet, wrapped around a large PEG molecule (Fig. S5-6). All six copies adopt essentially the same conformation and overlap closely, with rmsd=0.27 to 0.50 Å in pairwise overlaps of 10 Cα positions (residues 3-12). The conformation of the cyclic core region is very similar in all copies, characterized by a side-by-side stacking arrangement of D-Trp7 and Lys8, known to be important for SS receptor binding and activation (*28*) (Fig. S5-6). The D-Trp7-Lys8 pair of side chains is oriented slightly differently in copy D of the 6 molecules in asymmetric unit (Fig. S5) as a result of crystal-packing interactions (Fig. S5). Residues outside of the core (residues 1-4 and 11-13) show slightly more flexibility than those of the cyclic core itself.

**Fig. 4.**
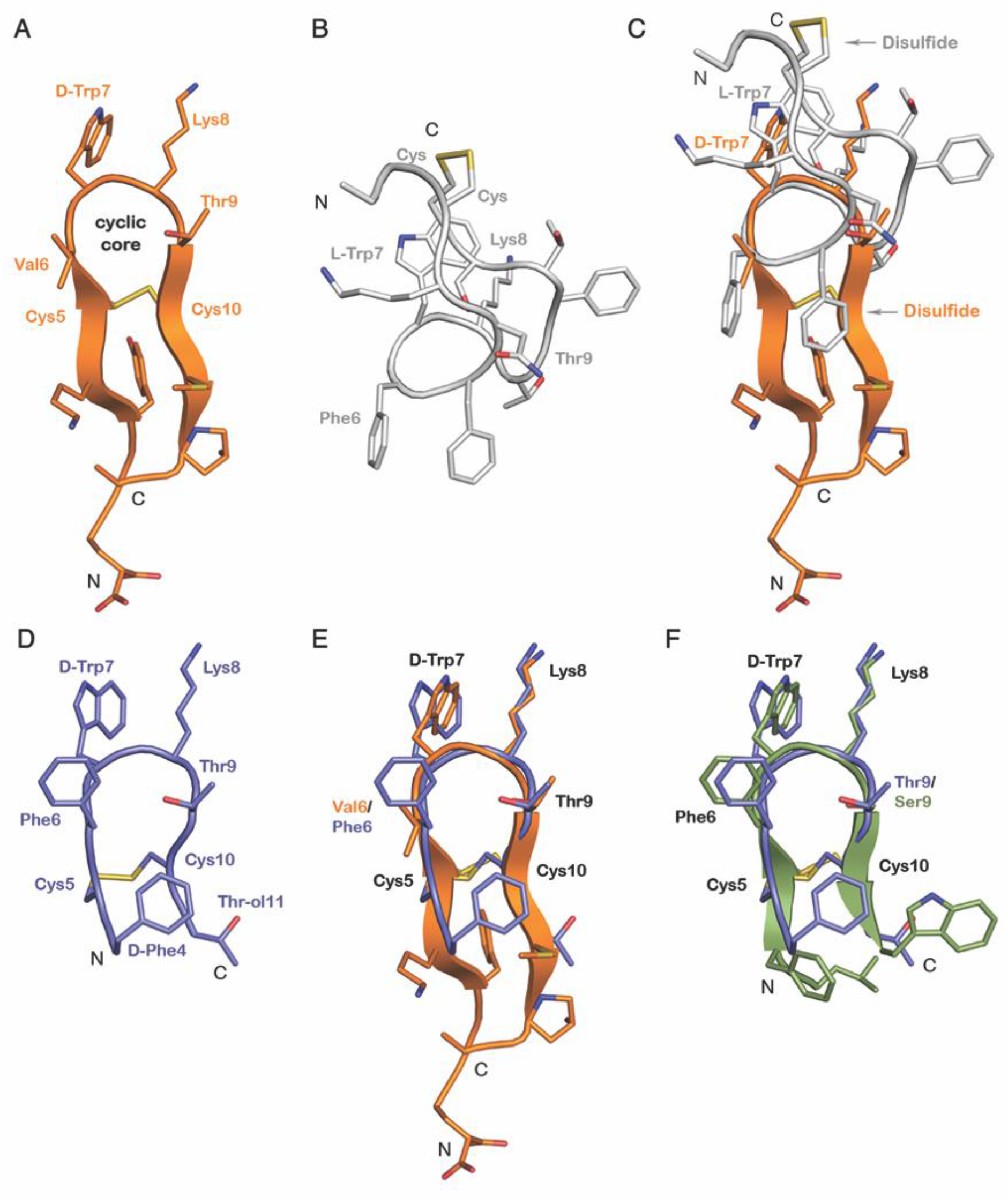
Consomatin Ro1 and G1 are structural mimetics of the SS drug analog Octreotide. **A.** X-ray structure of Consomatin Ro1 at 1.95 Å resolution. **B.** NMR solution structure of human SS-14 obtained during heparin-induced fibril formation (PDB ID: 2MI1). **C.** Alignment of the structure of Consomatin-Ro1 (orange) with that of SS-14 (gray) **D.** NMR solution structure of Octreotide (PDB ID: 1SOC) **E.** Alignment of the structure of Consomatin-Ro1 (orange) with that of Octreotide (purple) showing nearly identical backbone conformation and orientation of D-Trp7, Lys8, Thr9 and the disulfide bond, but differences in the amino acid composition and side chain arrangements of Val/Phe6 and of residues outside the cyclic core. **F.** Alignment of a homology model of Consomatin G1 (green), based on the structure of Ro1, with that of Octreotide (purple) suggesting that the molecules share strong structural similarities. Numbering of residues according to that of Consomatin Ro1.

Despite the sequence similarities between Consomatin Ro1 and human SS (Protein Data Bank (PDB) ID: 2MI1), the backbone conformations and side chain orientations of the two peptides are quite distinct (Fig. 4B-C). The apex of the turn of human SS “cups” in the opposite direction of Consomatin Ro1, making it impossible to overlap the structure on more than the 4 residues of the apex of the beta turn. Residues Trp7 and Lys8 of human SS (numbering taken from Consomatin Ro1) occupy similar positions although they are in different rotamers from Consomatin Ro1 (Fig. 4C).

In contrast, Consomatin Ro1 displays striking similarities with the solution structure of the SS drug analog Octreotide (PDB ID: 1SOC, Fig. 4D). The backbone of Consomatin Ro1 aligns closely with Octreotide, and their D-Trp7-Lys8-Thr9 residues adopt nearly identical conformations as in Octreotide (Fig. 4E). The most notable structural differences are seen in the side chain orientation of Phe6/Val6 within the cyclic core and between residues outside of the cyclic core motif (Fig. 4E). Thus, the venom peptide Consomatin Ro1 represents a nature-derived, structural mimetic of Octreotide and possibly other SS drug analogs for which structural data are not available.

Octreotide was developed as a stable drug agonist of the SST_2_, a receptor target for the treatment of neuroendocrine disorders such as acromegaly and gastroenteropancreatic neuroendocrine tumors (*39*). Correspondingly, Octreotide preferentially activated the SST_2_ over the SST_3_ and SST_5_ and displayed no significant activity on SST_1_ or the SST_4_ (*39*). These observations on the subtype selectivity of Octreotide and other SS analogs prompted us to investigate the activity of the Consomatin Ro1 venom peptide at the five human SSTs and related GPCRs.

### Consomatin Ro1 preferentially activates two of the five subtypes of the human SSTs

Consomatin Ro1 was screened against a panel of 318 human GPCRs, including all five subtypes of the SS receptor, using the PRESTO-Tango β-arrestin recruitment assay (*40*). When tested at 10 μM, Consomatin Ro1 activated the SST_4_ and, to a lesser extent, the SST_1_. At this concentration, the peptide did not activate any of the other 313 GPCRs tested, including the UTR and opioid receptors (DOR, KOR, MOR) (Fig. 5A). Subsequent concentration-response studies of Consomatin Ro1 at the five human SSTs confirmed preferential activation of the SST_1_/SST_4_ with little or no significant activity at the SST_2_, SST_3_ and SST_5_ (Fig. 5B). The EC_50_ value of Consomatin Ro1 is 2.9 μM (pEC_50_ ± CI95 = 5.5 ± 0.8) at the SST_1_ and 5.1 μM (pEC_50_ ± CI95 = 5.3 ± 0.3) at the SST_4_ (Fig. 5B, Table S4, calculated from 3 independent experiments each carried out in triplicate). EC_50_ values at the SST_2_, SST_3_ and SST_5_ could not be determined using a maximum concentration of 300 μM. For comparison, the EC_50_ of human SS-14 is between 1-97 nM for the five SSTs (Fig. 5, Table S4) which correlates well with previously reported receptor binding data (*41*). Consomatin Ro1, though having much lower potency than human SS, displays subtype preference towards the SST_1_/SST_4_. While not tested here, it is conceivable that the toxin is more potent at the homologous receptors expressed in fish which share ~ 60 % sequence identity with human SSTs (*29*).

**Fig. 5.**
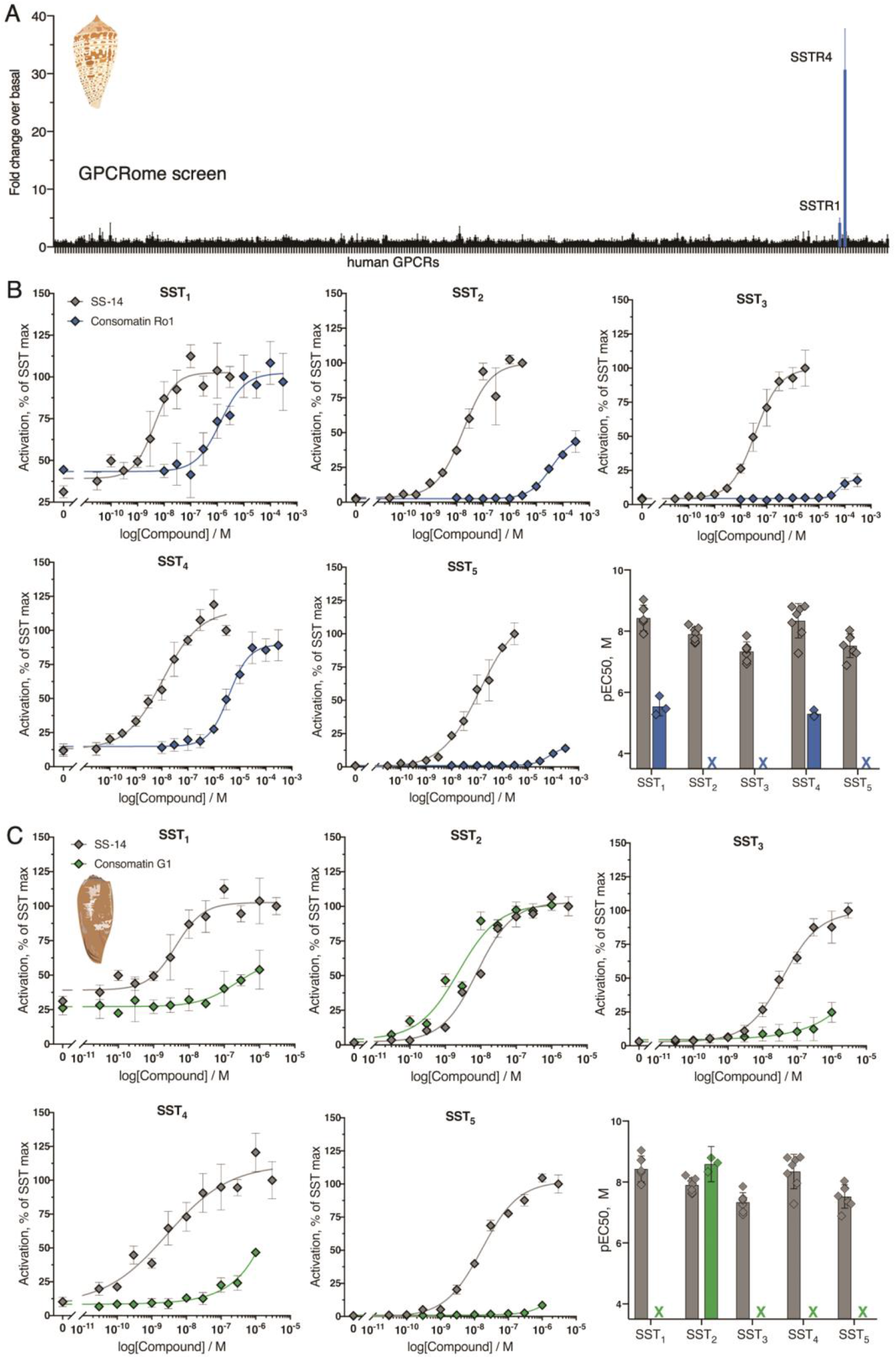
Consomatin Ro1 and G1 selectively activate the human SS receptors. **A.** PRESTO-Tango screen of Consomatin-Ro1 (10 μM) at 318 human GPCRs (x-axis) identified the SST_4_ and SST_1_ as the molecular target. Plotted values represent means ± SD. **B-C.** Representative concentration-response curves at the five human SS receptor subtypes comparing (**B.**) Consomatin Ro1 or (**C.**) Consomatin G1 with human SS-14 using the β-arrestin recruitment assay (error bars represent SD), as well as bar graphs showing the respective pEC50 values (error bars represent CI95). RLU = relative luminescent units.

### A single injection of Consomatin Ro1 provides antinociception and antihyperalgesia in two mouse models of acute pain

In addition to its inhibitory role in hormone secretion and cell proliferation, SS is expressed in sensory and sympathetic neurons where it can reduce nociception and inflammation (*42*). For example, the SS peptide mimetic, TT-232, that was initially developed for its endocrine antitumor activity (*43*), was later shown to provide analgesia in animal models of acute and neuropathic pain (*44*). Interestingly, the two subtypes that were associated with these effects were the SST_1_ and SST_4_ which are both activated by Consomatin Ro1 at low micromolar concentration. These observations led us to investigate the potential analgesic properties of the venom peptide using two mouse models of pain. In the natural setting, reducing the pain response in prey may help inhibit the escape response of fish after envenomation (*45*), thereby facilitating prey capture by slow-moving *Asprella* snails.

Antinociceptive effects were first evaluated through measurement of tail withdrawal latencies during noxious heat exposure using the tail flick test (*46*). In order to assess whether the toxin’s effects were mediated via receptors expressed in the central or peripheral nervous system, Consomatin Ro1 was either injected intrathecally (IT) or intraperitoneally (IP) prior to measurements. Mice injected IT with multiple doses of the venom peptide (0.025-0.75 mg/kg) did not show any significant increase in tail flick latencies compared to saline (Fig. 6A-B). Importantly, morphine (1 mg/kg, IT) injection (positive control) resulted in robust antinociception confirming the sensitivity of the assay (Fig. 6A-B). However, when Consomatin Ro1 was injected via the IP route, dose-dependent antinociception was observed, with the maximal effect observed at 2.5 mg/kg, that lasted up to 3 hours (Fig. 6C-D). Higher doses (5 and 7.5 mg/kg) did not result in any further antinociception. The observed effect was similar in magnitude to that of morphine at 3 mg/kg (Fig. 6C-D). It should be noted that the highest intrathecal dose tested (0.75 mg/kg) that demonstrated no efficacy was only 1/3^rd^ of the E_max_ of the IP dose. By contrast, typically 1/10-100^th^ of a systemic dose is effective when delivered IT, if a drug is acting in the spinal circuit (*47, 48*). Because Consomatin Ro1 is a peptide, it is highly unlikely that IP dosing resulted in crossing of sufficient amount of the drug across the blood brain barrier to induce an effect in the central nervous system (CNS). Together, these findings suggest that the analgesia-inducing site of action of Consomatin Ro1 is outside the CNS.

**Fig. 6.**
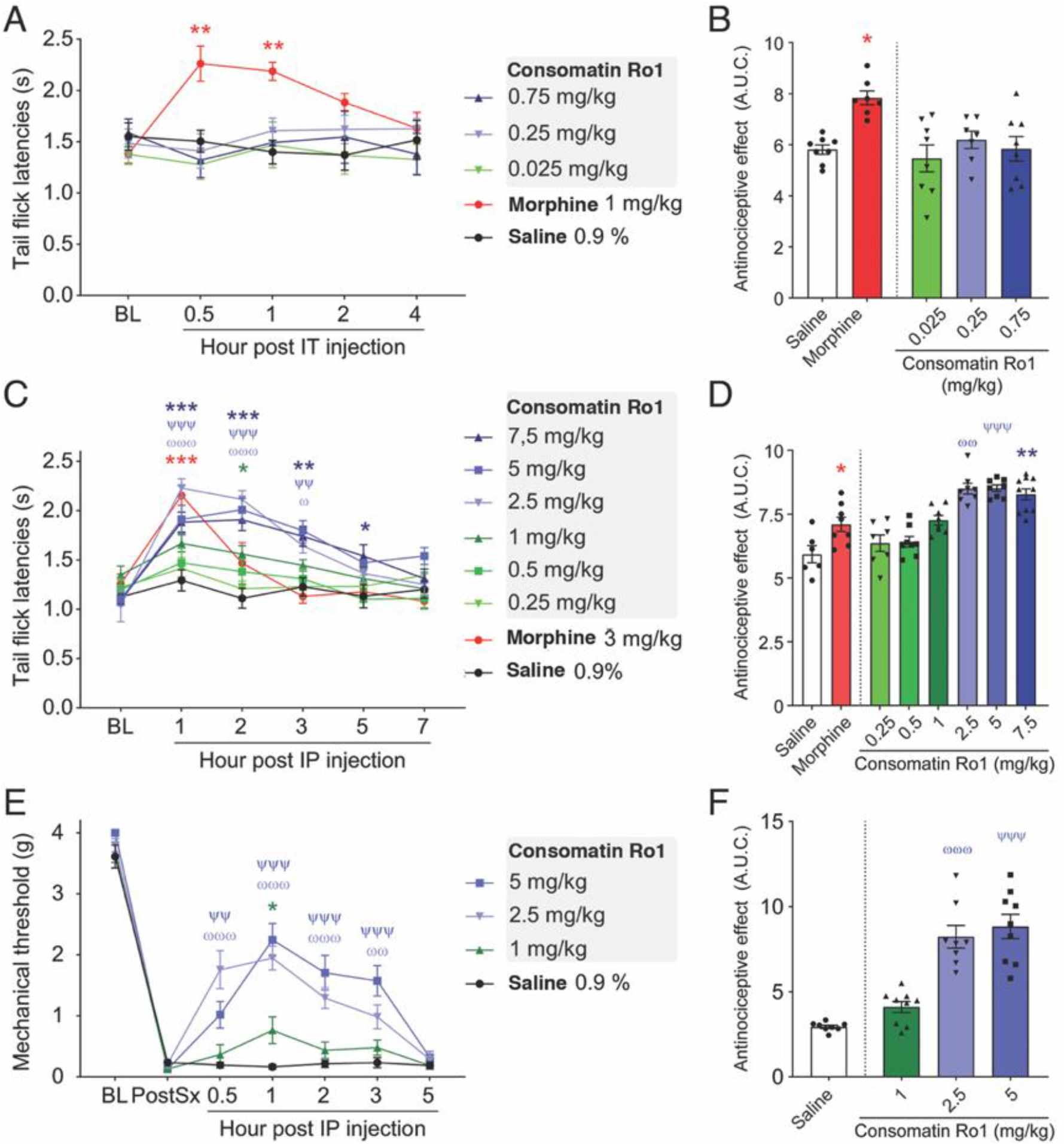
Consomatin Ro1 provides analgesia in two mouse models of acute pain. **A & B.** Sensitivity to thermal noxious stimuli was evaluated by dipping mouse tails into hot water (52°C) and recording tail flick latencies (TFLs). **A.** Mouse TFLs after a single intrathecal (IT) injection of saline, morphine, or multiple doses of consomatin-Ro1. TFLs were captured over a 4-hour period (n=7-8 per group, two-way ANOVA with Dunnett’s post-hoc test). **B.** Area under the curve between baseline and 4-hour condition, presenting global tail flick latencies when animals received IT injections (Kruskal-Wallis followed by Dunn’s post-hoc test). **C.** Mouse tail flick latencies after a single intraperitoneal (IP) injection of saline, morphine, or multiple doses of Consomatin Ro1. TFLs were captured over a 7-hour period (n=6-10 per group, two-way ANOVA with Dunnett’s post-hoc test). **D.** Area under the curve between baseline and 7-hour condition, presenting global tail flick latencies when animals received IP injections (Kruskal-Wallis followed by Dunn’s post-hoc test). **E.** Analgesic effect of Consomatin Ro1 on acute post-surgical pain. Plantar incision surgery was performed on the hind paw. 24-hours post-surgery, animals were administered saline solution or three increasing doses of Consomatin Ro1 through IP injections. Mechanical sensitivity was assessed using Von Frey filaments (n=8-9 per group, two-way ANOVA with Dunnett’s post-hoc test). **F.** Area under the curve between baseline and 5-hour condition, presenting mechanical withdrawal thresholds when animals received IP injections (Kruskal-Wallis followed by Dunn’s post-hoc test). Results corresponds to mean ± standard error of the mean (SEM). ^*/ω/ψ^p<0.05, ^**/ωω/ψψ^p<0.01, ^***/ωωω/ψψψ^p < 0.001, red asterisks illustrate significant difference between morphine and saline condition, blue asterisks: 7.5mg VS saline, ψ: 5mg VS saline ω: 2.5mg VS saline, green asterisks: 1 mg/kg VS saline.

We next evaluated the peptide’s efficacy in a mouse model of acute post-surgical pain (*49*). Following plantar incision surgery on the hind paw, mechanical sensitivity was determined by measuring the withdrawal response of the hind paw with a series of calibrated filaments (von Frey). 24 h after surgery, mice developed mechanical allodynia which did not change after IP injection of saline solution (control). However, a single injection of Consomatin Ro1 reversed mechanical hypersensitivity in a dose-dependent manner, with the effect lasting up to 3 hours (Fig. 6E-F). The maximal effect was observed at 2.5 mg/kg, similar to that observed in the tail flick assay. Similar effective IP doses have recently been reported for the small molecule SST_4_-agonist J-2156 in a rat model of breast cancer-induced bone pain (*50*). Together these data demonstrate that Consomatin Ro1 represents a new lead for the potential development of an analgesic that acts via opioid-independent pathways.

### Consomatin Ro1 defines a new class of venom evologs

The discovery of Consomatin Ro1 led us to search for other SS-like peptides in cone snail venom. In order to specifically identify candidates that likely activate the SS receptor, we limited our search to sequences that shared at least three of the four core residues important for receptor activation (Phe6-Trp7-Lys8-Thr9), or that had conservative substitutions at these positions. Sequence discovery was performed using the venom gland transcriptomes of *Asprella* snails and by assembling and mining the transcriptomes of 37 additional cone snail species available in the NCBI, DDBJ and the CNGB sequence repositories (Table S5). Searching a total of 602,377 assembled transcripts from these venom gland datasets led to the identification of 18 additional consomatins, two from the *Asprella* snails *C. rolani* and *C. neocostatus*, two from the net hunter *C. geographus*, and 14 from *Africonus* and *Varioconus*, two lineages of worm-hunting cone snails endemic to West Africa (*51*) (Fig. 7, Fig. S3). One of the *C. geographus* sequences was previously reported as “G042 Contulakin precursor conopeptide” (GenBank ID BAO65575) but its sequence similarity to SS was not recognized or stated (*52*). The name Contulakin was given based on the first C-superfamily toxin identified in *C. geographus*, Contulakin-G, a neurotensin-like peptide that activates the human neurotensin receptor (*53*).

**Fig. 7.**
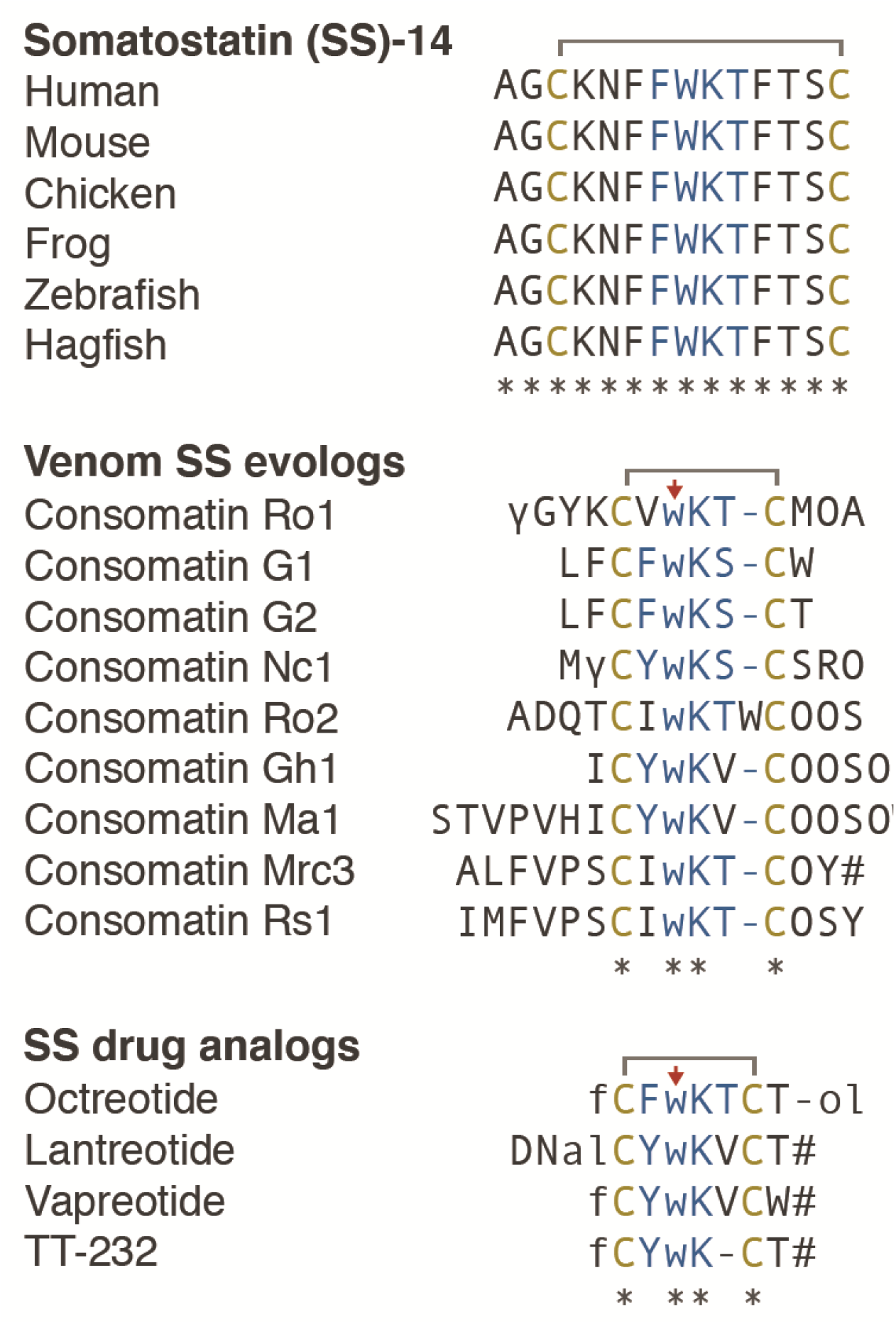
Comparative sequence alignments of vertebrate SS, selected consomatin evologs, and SS drug analogs. Identical amino acids are denoted by an asterisk (*). Mature toxins are shown with post-translational modifications based on processing and modification of Consomatin Ro1 (modifications and color codes as shown in Fig. 3). See Fig. S3 for precursor sequences of all consomatin sequences reported in this study.

The mature toxin regions of additional consomatins described here were predicted using the same proteolytic cleavage sites and post-translational modifications as those that appear to be involved in the processing of Consomatin Ro1. Sequence identity of these predicted mature toxins to human SS ranges from 14 % for Consomatin Ma1 to 38 % for Consomatin Ro1 (Fig. 7, Fig. S3). Notably, while SSs are highly conserved peptide hormones in vertebrates, sequences of consomatins are hypervariable with the exception of a conserved cysteine framework and Trp7-Lys8 motif (see sequence alignment in Fig. 7). The same pattern of variable and conserved residues can be observed for SS drug analogs, further emphasizing that consomatins represent a new, unexplored class of drug-like ligands of the SSTs or related receptors. To distinguish the venom peptides from the native prey hormones and synthetic hormone analogs we propose the name “evologs” for receptor ligands evolved by an organism to mimic the native ligand in another animal. Because of their streamlined role in inducing specific physiological changes in the other animal, evologs have distinctive biochemical and functional properties that make them ideal biomedical tools and drug leads.

Astonishingly, the sequences of the two consomatin evologs identified in the fish hunter *C. geographus*, Consomatins G1 and G2, are nearly identical to the drug Octreotide (80-90 % overall sequence similarity, Fig. 7). An alignment of a structural model of Consomatin G1 with the structure of Octreotide underscores their close similarities (Fig. 4F). Consistent with this observation, synthetic Consomatin G1 is a highly potent and selective activator of the human SST_2_ (EC_50_ = 2.6 nM, pEC_50_ ± CI95 = 8.6 ± 0.6, Fig. 5C, Table S4). Given that the SST_2_ is an important pharmacological target for the treatment of acromegaly and neuroendocrine tumors (NETs) (*39*), Consomatin G1 has potential as a therapeutic for the treatment of these disorders. In addition to its role in inhibiting GH secretion from the pituitary gland, SST_2_ is expressed in the pancreas where its activation suppresses the release of insulin and glucagon (*54*). Considering that Consomatin G1 was identified in the venom gland transcriptome of *C. geographus*, a species that uses fast-acting insulins to induce hypoglycemia in its fish prey (*16*), we hypothesize that Consomatin G1 may have evolved to modulate pancreatic hormone secretion, thereby exacerbating a hypoglycemic phenotype.

We note that the true chemical identity of Consomatin G1 and other consomatins identified from venom gland transcriptome data may differ from our predictions, particularly with respect to the nature of post-translational modifications and proteolytic N- and C-terminal cleavage sites. Thus, future studies on extracted venom are needed to elucidate the nature of the mature peptides deriving from these SS-like sequences. However, although the exact chemical identity and biological role of Consomatin G1 in the venom of *C. geographus* are yet to be experimentally determined, the discovery of additional SS-like peptides capable of activating the human SSTs at nanomolar potency establishes the evolution and wider use of SS evologs in cone snails and the presence of SS-like peptides in invertebrate deuterostomes in general. Furthermore, the presence of a large number of consomatins in several worm-hunting species suggests the existence of an SS-like receptor in marine worms with similar ligand binding sites to the vertebrate SSTs. Identifying such receptor-ligand system in the future will likely provide fundamental insight into the evolution of the SS signaling system.

Bioinformatic transcriptome mining revealed that several of the 39 analyzed cone snail species express C-superfamily sequences that, while not meeting our criteria for being SS-like, share a pair of cysteines and two of the four cyclic core residues with SS and related peptide hormones (Trp7-Lys8). Two of these sequences, Contulakin-Lt1 and Conlulakin-Lt2, from the venom gland transcriptome of *Conus litteratus* (Uniprot ID Q2I2P1 and Q2I2P2 (*55*)), were previously reported to have sequence similarity to an invertebrate UII-like peptide (*56*). Their biological activity was not assessed. Whether the *C. litteratus* toxins and other C-superfamily sequences with lower sequence similarity to SS evolved to target the SSTs, or another receptor family has not been addressed, but likely represents an untapped opportunity for the identification of additional GPCR ligands with interesting biological activities.

## Conclusions

Somatostatin is a critical hormone in health and disease. Decades of research have led to the design of metabolically stable, subtype-selective SS analogs for therapeutic application. Here, we show that through predator-prey evolution, venomous snails have mastered SS drug design to likely induce diverse physiological endpoints in prey. With hundreds of cone snail species yet to be sequenced, we anticipate the future discovery of many more consomatins with diverse receptor activation profiles that exert distinctive systemic effects. Together with our recent discovery of fast-acting venom insulins that has led to the design of a new drug lead for the treatment of diabetes (*24, 57*), this study provides a powerful example of the evolution of optimized, drug-like evologs of human hormones in venomous predators.

## Materials and Methods

### Specimen collection

*Conus rolani* and *Conus neocostatus* snails were collected from the waters of Sogod, Cebu, Philippines by tangle nets at depths of around 150-300 m. Collection of the snails was done under the Gratuitous Permits (GP-0053-11, 0063-12, GP-0084-15) issued by the Bureau of Fisheries and Aquatic Resources to the University of the Philippines Marine Science Institute. Specimen identification was initially performed by morphological examination and later verified by sequence analysis of the cytochrome oxidase c subunit 1 (COI) gene as previously described (*58*). For DNA/RNA analysis venom glands were dissected and stored in RNAlater (Thermo Fisher Scientific) until analysis. For venom extraction venom glands were dissected and stored at −80°C until analysis.

### Phylogenetic analysis

Phylogenetic analysis was performed using the *cytochrome* c oxidase subunit I (*COI*) barcode sequence for each species. Sequences were aligned and neighbor-joining trees generated using T-Coffee with default options (*59*). The tree was visualized using Tree Dyn 198.3 (*60*).

### Extraction and fractionation of *C. rolani* venom

Frozen venom glands of *C. rolani* were thawed in 30 % acetonitrile (CH_3_CN) with 0.2 % trifluoroacetic acid (TFA). Venom glands were cut with fine scissors and then homogenized. The mixture was centrifuged at 15,000 x rcf for 10 minutes as previously described (*61*). The supernatant was fractionated by reverse-phase (RP) HPLC on a Vydac preparative C_18_ column. Aliquots of the fractions and fraction pools were dried for bioactivity testing. Active HPLC fractions were further subfractionated using an analytical C_18_ column (Vydac 238TP; 5 μm, 250 x 4.60 mm). Gradient elution was done at a flow rate of 1 mL/min with increasing solvent B concentration (90 % CH_3_CN in 0.1 % TFA) in solvent A (nanopure water acidified with 0.1 % TFA) at 0.1 % Solvent B/min. Absorbance was measured at 220 nm and 280 nm.

### Mass determination and sequencing of Consomatin Ro1

The mass of the active fraction was determined using electrospray ionization mass spectrometry (ESI MS) at the Salk Institute for Biological Sciences Mass Spectrometry Core Facility. Prior to sequencing, an aliquot of the fraction was reduced using 26 μL of 50 mM dithiothreitol (Sigma-Aldrich) and incubated at 65°C for 30 minutes. The reduced peptide was alkylated with 35 μL of 1.5 M iodoacetamide (Sigma-Aldrich) and incubated in the dark at room temperature for 15 minutes. All solutions were buffered with Tris (pH 7-8). The reduced and alkylated peptide was run in the HPLC from 10 % to 25 % Solvent B (90 % CH_3_CN in 0.1 % TFA) in 45 minutes. The amino acid sequence of the peptide was determined by automated Edman degradation at the Protein Facility of the Iowa State University and by tandem MS sequencing.

For tandem MS, an aliquot of the sample was dried in the speedvac and subsequently reduced and alkylated in vapor using 1 % 2-methylaziridine and 2 % trimethylphosphine in 50 % acetonitrile and 100 mM ammonium bicarbonate (pH 8.4) for 60 min at RT. Alkylation vapor was removed and the sample was reconstituted in 0.5 % acetic acid. 1 μg aliquots of the now reduced and alkylated sample were loaded onto a 75 μm × 50 cm Pepmap EasySpray column using the autosampler of an Easy nLC-1000 nano-HPLC coupled to an Orbitrap Fusion Lumos Tribrid mass spectrometer. The sample was eluted using a flow rate of 0.2 μL/min with a gradient of 0–100 % B (solvent A = 0.5 % acetic acid, solvent B = 90 % acetonitrile in 0.5 % acetic acid (*v*/*v*)) in 90 min, a spray voltage of 2.0 kV, and 3 sec cycle time. MS1 scans were acquired at 120,000 resolution (@ 400 *m/z*) 50 ms maximum injection time. MS2 was acquired on precursors that carry 4-20 charges using the following settings: 3 microscans, 2 *m/z* isolation window, target value of 5e4 ions, 200 ms maximum injection time. Each precursor was subjected to ETD and HCD fragmentation using the following conditions: 30,000 resolution (@ 400 *m*/*z*), ETD using 60 ms ion reaction time, HCD using 32 % normalized collision energy. Raw files were searched using search engine Byonic (Protein Metrics) allowing for a 10 ppm mass tolerance and common conotoxin PTMs. The peptide sequence was then manually verified.

### Transcriptome sequencing of the *C. rolani* and *C. neocostatus* venom gland

Total RNA was extracted from the venom glands of *C. rolani* and *C. neocostatus* using the Direct-zol RNA extraction kit (Zymo Research), with on-column DNase treatment, according to the manufacturer’s instructions. cDNA library preparation and sequencing were performed by the University of Utah High Throughput Genomics Core Facility as previously described (*24*). Briefly, 125 cycle paired-end sequencing was performed on an Illumina HiSeq2000 instrument at an 80 % standard cluster density. Adapter trimming of de-multiplexed raw reads was performed using fqtrim (v0.9.4), followed by quality trimming and filtering using prinseq-lite (*62*). Error correction was performed using the BBnorm ecc tool, part of the BBtools package (open source software, Joint Genome Institute). Trimmed and error-corrected reads were assembled using Trinity (version 2.2.1) (*63*) with a k-mer length of 31 and a minimum k-mer coverage of 10. Assembled transcripts were annotated using a blastx search (*64*) (E-value setting of 1e-3) against a combined database derived from UniProt, and an in-house cone snail venom transcript library.

### Peptide synthesis, folding and purification

Peptide synthesis. Consomatin Ro1 was synthesized using an Apex 396 automated peptide synthesizer (AAPPTec; Louisville, KY) applying standard solid-phase Fmoc (9-fluorenylmethyloxy-carbonyl) protocols. Peptide was constructed on preloaded Fmoc-Ala-Wang resin (substitution: 0.38 mmol/g; Peptides International, Louisville, KY). All standard amino acids were purchased from AAPPTec except for: N-α-Fmoc-O-t-butyl-L-trans-4-hydroxyproline (Hyp) was purchased from EMD Millipore (Billerica, MA), Fmoc-γ-carboxy-glutamic acid γ,γ-di-t-butyl ester (Gla) was purchased from Advanced ChemTech (Louisville, KY) and *N*_α_-Fmoc-*N*_(in)_-Boc-D-tryptophan (D-Trp) from Chem-Impex International, Inc (Wood Dale, IL). Side-chain protection for the amino acids was as follows: Cys, trityl (Trt); Gla, t-butyl ester (OtBu); Lys and D-Trp, tert-butyloxycarbonyl (Boc); Hyp, Thr and Tyr, tert-butyl ether (tBu). Tenfold excess of standard amino acids was used except for Fmoc-Gla(OtBu)_2_, Fmoc-Hyp(tBu) and Fmoc-D-Trp(Boc)-OH, for which 5-fold excess was used. The coupling activation was achieved with one equivalent of 0.4 M benzotriazol-1-yl-oxytripyrrolidinophosphonium hexafluorophosphate (PyBOP) and two equivalents of 2 M *N,N*-diisopropylethyl amine (DIPEA) in N-methyl-2 pyrrolidone (NMP) as the solvent. Each coupling reaction was conducted for 60 min except for the special amino acids for which the reaction was conducted for 90 min. Fmoc deprotection was carried out for 20 min with 20 % piperidine in dimethylformamide (DMF).

Peptide cleavage and purification. Consomatin Ro1 was cleaved from 103 mg resin by a 1.5-h treatment with 1 mL of Reagent K (TFA/water/ phenol/ thioanisole/1,2-ethanedithiol 82.5/5/5/5/2.5 by volume). Next, the cleavage mixture was filtered and precipitated with 10 mL of cold methyl-tert-butyl ether (MTBE). The crude peptide was then precipitated by centrifugation at 7,000 x rcf for 6 min and washed once with 10 mL cold MTBE. The crude peptide was purified by reversed-phase HPLC using a semi-preparative C_18_ Vydac column (218TP510, 250 x10 mm, 5-μm particle size) eluted with a linear gradient ranging from 10 % to 40 % solvent B in 30 min at a flow rate 4 mL/min. The HPLC solvents were 0.1 % (vol/vol) TFA in water (solvent A) and 0.1 % TFA (vol/vol) in 90 % aqueous acetonitrile (vol/vol) (solvent B). The eluent was monitored by measuring absorbance at 220/280 nm. Purity of the peptide was assessed by analytical C_18_ Vydac RP-HPLC (218TP54, 250 x 4.6 mm, 5-μm particle size) using the same gradient as described above with a flow rate 1 mL/min. The peptide was quantified by UV absorbance at 280 nm, using an extinction coefficient (ε) value of 6990 M^-1^·cm^-1^. Out of 103 mg of the cleaved resin, 8.3 mg of the linear peptide was prepared.

Oxidative folding of Consomatin Ro1. Two hundred nanomoles of linear Consomatin Ro1 was resuspended in 1 mL of 0.01 % TFA solution and added to a solution containing: 5 mL of 0.2 M Tris·HCl (pH 7.5) plus 0.2 mM EDTA, 0.5 mL of 20 mM reduced and 0.5 mL of 20 mM oxidized glutathione, and 3 mL of water. Final peptide concentration in the folding mixture was 20 μM. The folding reaction was conducted for 2 h and quenched with formic acid to a final concentration of 8 %. The quenched reaction mixture was then separated by RP-HPLC using a semi-preparative C_18_ column and a linear gradient ranging from 10 % to 40 % of solvent B in 30 min with a flow rate of 4 mL/min. The eluent was monitored by absorbance at 220/280 nm. Purity of the folded peptide was assessed by an analytical C_18_ Vydac RP-HPLC using the gradient described above, with a flow rate 1 mL/min. Pure fully folded Consomatin Ro1 was quantified by absorbance at 280 nm as described for the linear peptide, which was later verified by the Amino Acid Analysis performed by the University of Utah DNA/Peptide Facility. 0.71 mg of the final Consomatin Ro1 was obtained from 1.89 mg of the linear peptide with 98 % purity. The molecular mass of Consomatin Ro1 was confirmed by ESI MS (calculated monoisotopic MH^+1^:1573.64, determined monoisotopic MH^+1^: 1573.64), at the University of Utah Mass Spectrometry and Proteomics Core Facility. An additional mass of 1595.63 Da was also identified, which corresponded to sodium adduct: [M+Na]^+^ ion.

Consomatin G1 were custom synthesized by GenScript. The peptides’ mass and purity (>95 %) were verified by RP-HPLC and MS.

### Coelution of native and synthetic peptides

The native and synthetic Consomatin Ro1 were separately loaded on an analytical C_18_ column (Vydac 238TP; 5 μm, 250 x 4.60 mm) and profiled using a linear gradient from 19 % to 21 % Solvent B in 20 minutes. A 1:2 mixture of the native and synthetic Consomatin Ro1 was also applied on the same column using the same gradient and conditions.

### Determination of the crystal structure of Consomatin Ro1

Lyophilized Consomatin Ro1 was dissolved in water at 40 mg/mL and filtered with a 0.2 μm filter. This solution was diluted with water for crystallization trials. For crystal screening, WT Consomatin Ro1 (at 20 mg/ml) was mixed in a 1:1 ratio with commercial crystallization solutions in sitting drop vapor diffusion crystal trays. Consomatin Ro1crystallized after 11 days at 21 °C in SaltRx (Hampton Research), condition D10 (4.0 M Sodium Nitrate, 0.1 M Sodium Acetate Trihydrate, pH 4.6). A WT crystal was prepared for data collection by suspending the crystal in a small nylon loop, briefly immersing in fresh mother liquor for 20 s, and cryocooling by plunging in liquid nitrogen. A Platinum heavy-atom-soaked crystal was prepared by soaking for 5 minutes in 20 μl fresh mother liquor containing 0.1 mM K_2_PtCl_4_. The crystal was then cryocooled by plunging in liquid nitrogen (without back soaking).

X-ray diffraction data were collected at the Stanford Synchrotron Radiation Lightsource (SSRL) with the crystal maintained at 100 K. WT data were collected with 1.7711Å X-rays in an effort to optimize sulfur anomalous scattering. Data were integrated and scaled using XDS (*65*) and AIMLESS (*66*) (Table 1). A fluorescence scan of a Platinum-soaked crystal indicated the presence of Platinum and the X-ray energy was chosen based on this scan to optimize anomalous scattering. The resulting Platinum derivative data were sufficient to determine initial phases by Single Wavelength Anomalous Diffraction (SAD) analysis using the program Autosol (*67*) in the PHENIX software package (*67*). The resulting electron density map was readily interpretable and an initial model was built using the program Coot (*68*). The initial model comprised most of 6 copies of Consomatin Ro1 and was partially refined (*69*) against the Platinum derivative data set. The model was then used to calculate model-based phases and the model-building and refinement (using CCP4 program Refmac5) (*70, 71*) continued using the WT data set. The model was initially refined to Rcryst= 0.275 and Rfree= 0.302 with good geometry. The model was evaluated using Molprobity (*72*) and judged to be of good quality, however, a large region of unexplained density was visible in the FoFc electron density map. The density strongly resembled that of a Polyethylene Glycol molecule, although PEG was not a component of the crystallization solution and was not thought to be present in the protein solution. Models of several PEG molecules were tested by model building and refinement until the best fit to the density was obtained with a molecule of composition C_33_O_17_H_67_ (predicted molecular weight ~750 Daltons). The model was improved by including the PEG molecule and was refined to Rcryst= 0.239 and Rfree= 0.283 with good geometry.

The structural homology model of the venom peptide Consomatin G1 was predicted by changing the side chains of Ro1 to those of Consomatin G1 in Coot (*68*), based on the sequence alignment (Fig. 3).

### Plasma stability assays

The *in vitro* stability of Consomatin Ro1 was investigated in human plasma (mixed gender) and compared to human somatostatin-14. Peptides were incubated in 500 μL plasma at 37 °C with a final concentration of 3 μM. Samples (40 μL) were taken at 0, 5, 15, 30, 60, 120, 240, 480, and 1440 min time points. A 2-fold volume of cold acetonitrile with 0.1 % formic acid was added to terminate the incubation. Samples were applied on a C_18_ column (Waters CSH, 2.1 × 50 mm, 1.7 μm particle size) with a flow rate of 0.650 mL/min and a gradient profile of 2 % to 20 % Solvent B in 3.5 min and then 20 % to 98 % Solvent B in 0.5 min and run on a Thermo Vanquish Horizon UHPLC. The HPLC solvents were 0.1 % formic acid in water (solvent A) and acetonitrile (solvent B). MS data was acquired using a Thermo Q-Exactive Focus Orbitrap mass spectrometer at 35,000 resolution (FWHM @ m/z 200) for full scan and 17,500 resolution for MS2 in DDI mode. Parent compound disappearance, based on relative LC/MS peak area (0 min = 100 %), was used to calculate half-lives. Propanthelin bromide (1 μM) was used as a control and two replicates for each compound were analyzed.

### Initial GPCRome Screening

Functional data were generously provided by the National Institute of Mental Health’s Psychoactive Drug Screening Program, Contract # HHSN-271-2013-00017-C (NIMH PDSP). The NIMH PDSP is Directed by Bryan L. Roth at the University of North Carolina at Chapel Hill and Project Officer Jamie Driscoll at NIMH, Bethesda MD, USA. Consomatin Ro1 was tested on a panel of GPCRs using the PRESTO-Tango system (*40*) which uses HTLA cells with a tTA-dependent luciferase reporter and a β-arrestin2 TEV protease fusion gene. HTLA cells were plated in 384-well plates overnight before transfection of the receptor constructs (318 target receptors). Assay plates with the compounds were incubated overnight at 37 °C. The compounds and the media were removed and Bright-Glo reagent (Promega) was added to determine luciferase activity. Results were presented in fold of the average basal; normal activity is 0.5-to 2.0-fold of basal level (~4 RLU). Dopamine receptor DRD2 with 100 nM quinpirole was used as an assay control. Follow-up assays were done on the somatostatin receptors.

### β-arrestin recruitment assay (PRESTO-Tango)

HTLA cells (a kind gift from Bryan L. Roth) were maintained in DMEM (ThermoFisher) supplemented with 10 % FBS (Biowest), 100 U/mL penicillin and 100 μg/mL streptomycin (ThermoFisher) (HTLA medium) with 100 μg/mL Hygromycin B (ThermoFisher), and 2 μg/mL puromycin (ThermoFisher) added. On day 1, 3 million cells were seeded in t25 flasks in 5 mL HTLA medium and incubated overnight. On day 2, medium was changed, and cells were transfected using Polyfect (Qiagen) according to manufacturer’s protocol. On day 3, the cells were resuspended in DMEM supplemented with 1 % dFBS (ThermoFisher) (assay medium), and 15,000 cells in 40 μL per well were seeded in poly-D-lysine-coated white clear-bottom 384 well plates (Corning) and incubated overnight. On day 4, medium was changed, and ligands were diluted to appropriate concentrations (5 x final concentration) in HBSS (Thermofisher) supplemented with 20 mM HEPES (Sigma), 1 mM of CaCl_2_, 1 mM of MgCl_2_, pH adjusted to 7.4 with 10 M NaOH (assay buffer), which was supplemented with 0.1 % BSA (ligand buffer). 10 μL/well was added, and cells were incubated overnight. On day 5, the medium and compounds were removed from the cells, and 20 μL/well of a 1:20 dilution of BrightGlo (Promega,) in assay buffer supplemented with 0.01 % pluronic F68 (ThermoFisher) was added to the cells. Plates were incubated for 20 minutes in the dark at room temperature, and luminescence was measured on a Molecular Devices SpectraMax iD5 (Molecular Devices) with each well integrated for 1 second.

### IACUC Approval

All animal studies were approved by the Institutional animal care and use committee (IACUC) at the Universities of Utah and Arizona. Protocol numbers: #14-08018 (approved period, August 20, 2014 - August 19, 2017), #17-07020 (approved period, August 01, 2017 - July 31, 2020), and #15-578 (approved period, July 3, 2018 - July 3, 2021).

### Intracranial mouse bioassay

Swiss Webster mice (12 to 22 days old) were injected intracranially (i.c.) with the peptide dissolved in 15 μL Normal Saline Solution (NSS; 0.9 % NaCl). Mouse behaviors in response to stimuli such as prodding were observed for at least an hour. Behavioral differences between the treated and control groups were recorded.

### Intraperitoneal (IP) administration of Consomatin Ro1

For intraperitoneal injection, the injection site (left lower abdominal quadrant) was wiped with 2% chlorhexidine. Mice were manually restrained, abdomen side up, with cranial end pointed down. With an angle of 15 to 20 degrees, the abdominal cavity was punctured to inject either 200 μL of saline solution, 0.25, 0.5, 1, 2.5, 5 or 7.5 mg/kg of Consomatin Ro1. A solution of morphine sulfate (3 mg/kg) was injected as positive control. Before injection, aspiration was attempted to ensure that an abdominal viscus had not been penetrated.

### Intrathecal (IT) administration of Consomatin Ro1 by lumbar puncture

For single administration, mice were sedated with 2% isoflurane in O_2_ anesthesia delivered at 2 L/min prior to the injection. Increasing doses (0.025 to 0.75 mg/kg) of Consomatin Ro1 or morphine (1 mg/kg) were administered by lumbar puncture as described by Hylden and Wilcox (*73*). Briefly, mice were lightly restrained by the pelvic girdle in one hand (shaved, skin wiped with 2% chlorhexidine), while the syringe was held in the other hand at an angle of about 20° above the vertebral column. A 21G needle was inserted into the tissue to one side of the L5 or L6 spinous process so that it slipped into the groove between the spinous and transverse processes. The needle was then moved carefully forward to the intervertebral space as the angle of the syringe was decreased to about 10°. The tip of the needle was inserted so that approx. 0.5 cm was within the vertebral column. The solution was then injected in a volume of 5 μl and the needle rotated on withdrawal.

### Measurement of thermal sensory thresholds – Tail flick test

Antinociceptive effect was assessed through measurement of tail withdrawal latencies using the tail flick test. A hot water chamber was set at 52°C. Adult mice were restrained in a cylinder and the distal 2/3 of the tail was dipped into hot water. To record tail flick latencies, a chronometer was started when the mouse tail was dipped into water and stopped as soon as the mouse withdrew its tail. Results correspondsto the mean time when animals start to flick their tail out of the water. A cut off time is set at 10 seconds before manual removal of the tail to prevent tissue damage.

### Paw incision model of post-surgical pain

To analyze acute post-surgical pain, animals were subjected to an incisional surgery based on the procedure previously described by Brennan et al. (*49*). A 0.5 cm long incision was made through skin and fascia of the plantar surface of the left hind paw. The plantaris muscle was then elevated and longitudinally incised, leaving the muscle origin and insertion intact. After hemostasis with gentle pressure, the skin was closed with two mattress sutures of 5-0 nylon on a curved needle. Sham animals underwent anesthesia, and their left hind paw was scrubbed with betadine and ethanol (70%). Only incision was performed, without damaging the muscle.

### Measurement of tactile sensory thresholds – Von Frey test

The assessment of tactile sensory thresholds was determined by measuring the withdrawal response to probing the plantar surface of the hind paw with a series of calibrated filaments (von Frey) 24 hours post surgery. Each filament was applied perpendicularly to the plantar surface of the paw of a mouse held in suspended wire mesh cages. The ‘‘up and down’’ method was used to identify the mechanical force required for a paw withdrawal. Data were analyzed with the nonparametric method of Dixon, as described by Chaplan et al. (*74*). Results are expressed as the mean withdrawal threshold that induces a paw withdrawal response in 50% of animals.

## Acknowledgements

We thank Judit Erchegyi, Charleen Miller, and our friend and colleague Jean Rivier (†2019) for peptide synthesis, Kevin Chase for assistance with transcriptome assemblies, Peter Huynh and Samuel D. Robinson for initial behavioral recording, Neda Barghi and Arturo O. Lluisma for help with initial sequence analysis, Paula Flórez-Salcedo for shell illustrations, Noel Saguil for assistance with specimen collection under the Department of Agriculture – Bureau of Fisheries and Aquatic Resources-issued gratuitous permit no. 0053-11, 0063-12, and 0084-15. We also thank the NIMH Psychoactive Drug Screening Program for initial screening of Consomatin Ro1, the High Throughput Genomics Core Facility, the DNA Sequencing Core Facility, the DNA/Peptide Facility, and the Mass Spectrometry and Proteomics Core Facility at the University of Utah (mass spectrometry equipment was obtained through a Shared Instrumentation Grant 1 S10 OD018210 01A1), the Iowa State University for Edman sequencing, and William Low at the Salk Institute for initial mass spectrometric analysis.

The mass spectrometric experiments were in part supported by a shared instrumentation grant from the NIH, 1S10OD010582-01A1, for the purchase of an Orbitrap Fusion Lumos Tribrid mass spectrometer. Use of the Stanford Synchrotron Radiation Lightsource, SLAC National Accelerator Laboratory, was supported by the U.S. Department of Energy, Office of Science, Office of Basic Energy Sciences under Contract No. DE-AC02-76SF00515. The SSRL Structural Molecular Biology Program is supported by the DOE Office of Biological and Environmental Research, and by the National Institutes of Health, National Institute of General Medical Sciences (P41GM103393).

## Funding

This research was funded by a Department of Defense Grant (W81XWH-17-1-0413 to B.M.O.), a Villum Young Investigator Grant (19063 to H.S-H.), a Department of Science and Technology – Philippine Council for Health Research and Development grant FP 140015 (to G.P.C.), a USAID/Philippines through the Science, Technology, Research, and Innovation for Development (STRIDE) Scholarship Program grant (to I.B.L.R.).

## Author contributions

I.B.L.R. and H.S.-H. discovered Consomatin Ro1, I.B.L.R. performed bioactivity testing in mice, I.B.L.R. and J.S.I. carried out venom extractions, fractionations and compound purifications. J.G. synthesized peptides, M.W. performed phylogenetic analysis, D.T. maintained cone snails and performed behavioral observations and recording, I.B.L.R., H.S.-H., W.R. performed venom extractions, derivatization and mass spectrometric sequencing, B.U. led mass spectrometric sequencing, I.B.L.R. and W.B.-Y performed and analyzed GPCR assays, W.B.-Y and H.B.-O. led GPCR assays, I.B.L.R. crystallized proteins, F.G.W. performed crystallographic analyses, C.P.H. oversaw X-ray crystallography, L.F.M. performed pain assays, A.P. led pain assays, H.S.-H. performed transcriptomics analyses, G.P.C., B.M.O. and H.S.-H. led the research. H.S.-H. wrote the manuscript.

## Competing interests

The authors declare no competing interests

## Data and materials availability

Structural data of Consomatin Ro1 will be available in the PDB (accession number pending). Sequences of Consomatins identified here are available in GenBank under the following accession numbers: Consomatin Ro1: MT409178, Consomatin Ro2: MW390882, Consomatin Nc1: MW390884, Consomatin G1: MW390883, Consomatin G2 (annotated as G042 Contulakin precursor conopeptide): BAO65575.

## Supplementary Materials

Materials and Methods

Table S1-S5

Figs S1-S6

Movies S1-S6

## Notes

### Competing Interest Statement

The authors have declared no competing interest.

