## Supplementary data (figures S1-S6, tables S1-S5) for "Somatostatin venom analogs evolved by fish-hunting cone snails: From prey capture behavior to identifying drug leads"

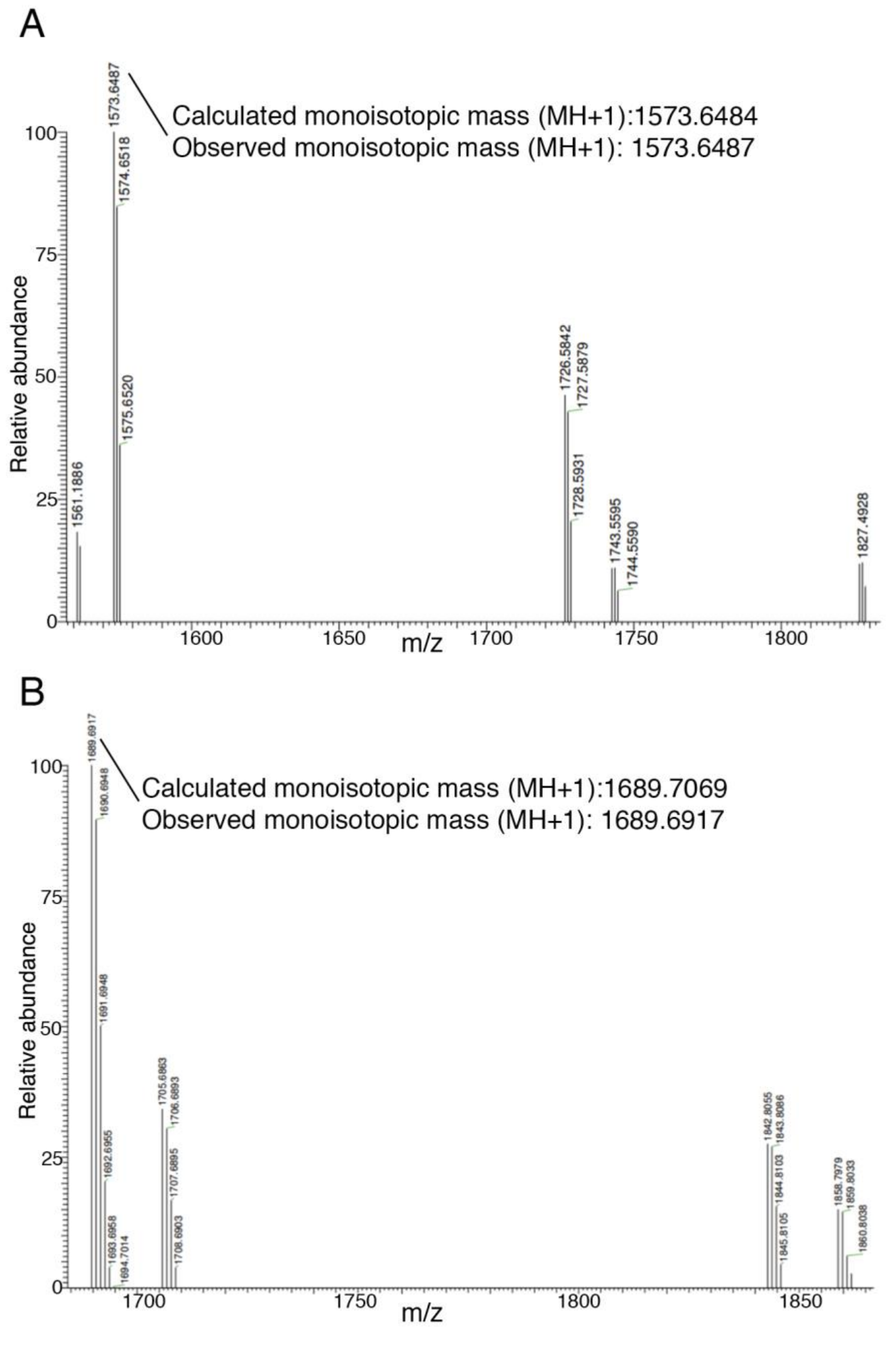


**Fig. S1.** Mass determination of compounds in fraction #16-12 confirming the molecular mass of Consomatin Ro1 sequenced by Edman and transcriptome analysis. **A.** Nonderivatized fraction 16-12. **B.** Fraction derivatized with dithiothreitol (DTT) and iodoacetamide (IAA).

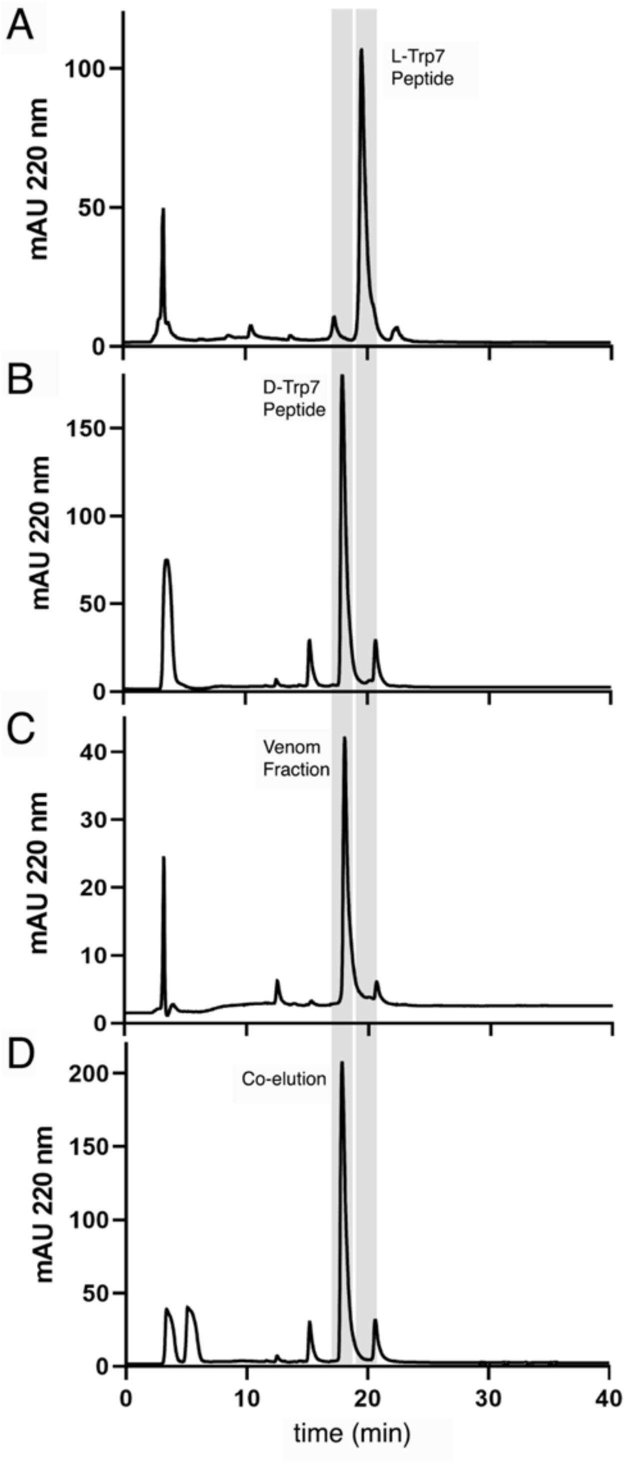


**Fig. S2.** Reverse-phase elution profile of **A**. Consomatin Ro1 containing L-Trp7, **B**. Consomatin Ro1 containing D-Trp7, **C**. the venom fraction #16-12, and **D**. co-elution of Consomatin Ro1 (D-Trp7) with the venom fraction.

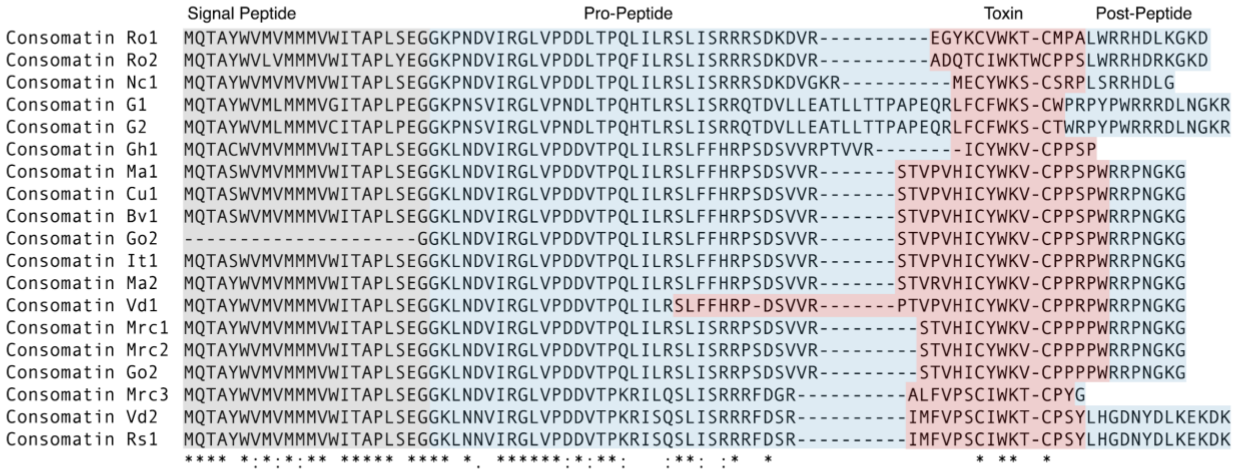


**Fig. S3.** Alignment of the precursor sequences of Consomatin Ro1 and other consomatins identified in the venom gland transcriptome of *Conus rolani* (Ro), *Conus neocostatus* (Nc), *Conus geographus* (G), *Conus grahami* (Gh), *Conus maioensis* (Ma), *Conus cuneolus* (Cu), *Conus boavistensis* (Bv), *Conus galeao* (Go), *Conus infinitus* (It), *Conus verdensis* (Vd), *Conus mercator* (Mrc), *Conus raulsilvai* (Rs1). Precursor regions encoding the predicted mature toxins are shown in red (signal peptide in gray and pro-and post-peptides in blue). Amino acids with sequence similarity to SS are shown in bold. Identical amino acids are denoted by an asterisk (*). Full stops (.) and colons (:) represent a low and high degree of similarity, respectively. Names of toxins that were synthesized and functionally characterized here are shown in bold. SRA accession numbers are provided in Table S4.

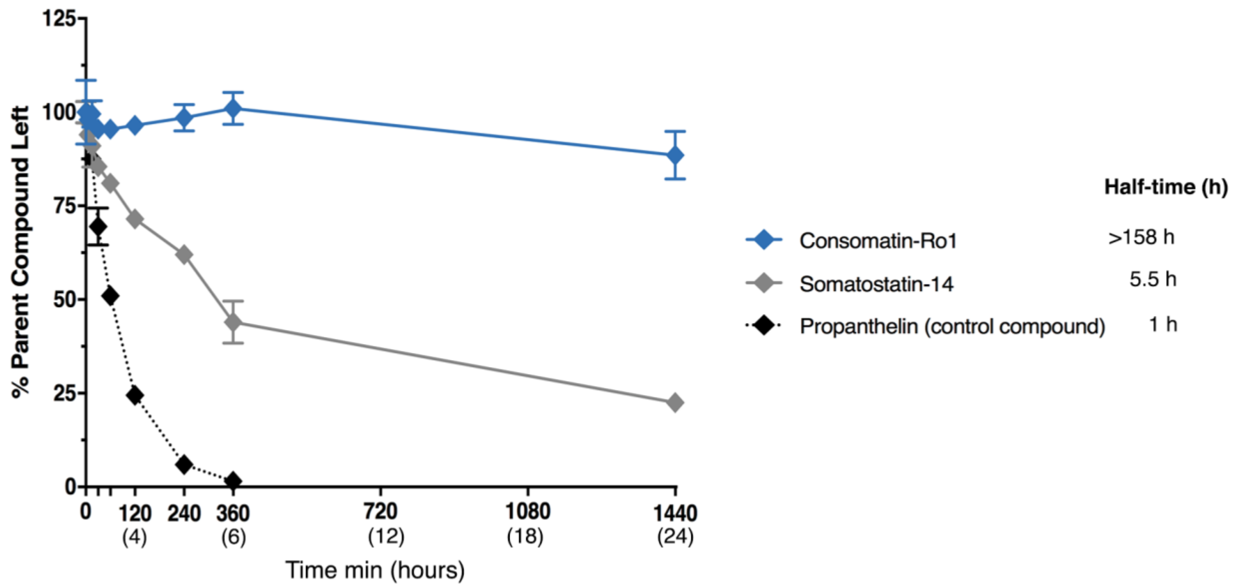


**Fig. S4.** Stability profiles of human SS-14, Consomatin Ro1, and an assay control compound, Porpenthelin, measured over 24 hours. Parent compound disappearance was based on relative LC/MS peak area (0 min = 100%). The values represent the mean ± S.D. of duplicate samples.

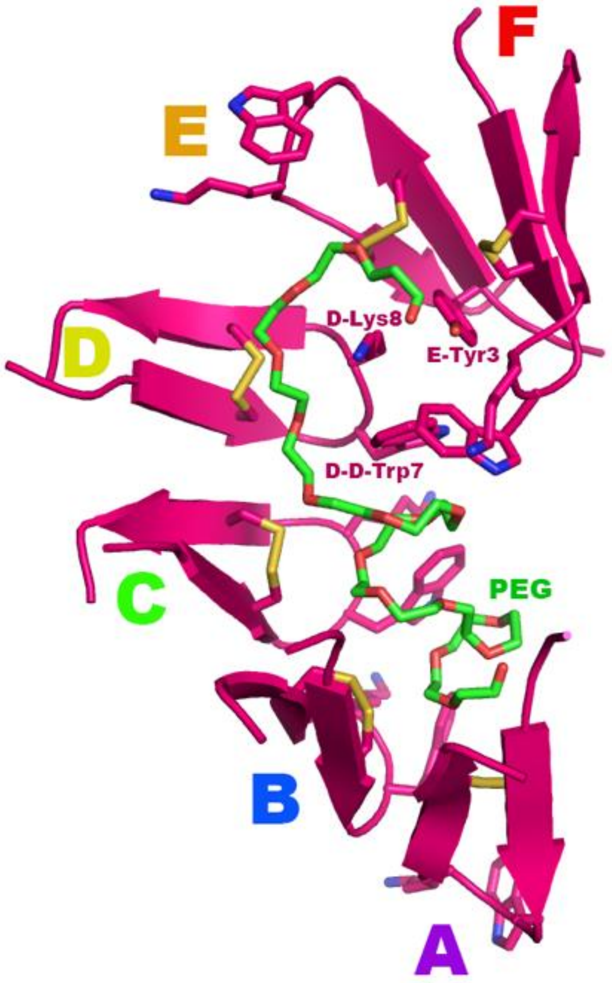


**Fig. S5.** Packing of molecules in the crystal. Six Consomatin Ro1 molecules (pink ribbons), and one large PEG molecule (green sticks with red oxygen atoms) form the asymmetric unit in the crystal. The origin of the PEG molecule in the crystal is not apparent, but it provided a good description of the electron density. Chain identifiers, A,B,C,D,E,F, are spectrum-coded to match the overlapped models in figure S6. The C5-C10 Cys-Cys disulfide bond of each molecule is shown as sticks (yellow). The two residues at the apex of the cyclic core (cysteine loop) (D-Trp7 and Lys8) are shown as sticks with blue nitrogen atoms. The six copies of Ro1 form a curving anti-parallel beta sheet with the apex of the turn of each copy of Ro1 oriented to the same side of the sheet except for copy E which is oriented opposite the others. Although copy E is oriented opposite the others, the same "top side" of the molecule faces the inner curve of the arc where the disulfide bond of all six molecules interacts with the PEG molecule. The PEG molecule facilitates crystallization primarily through hydrophobic interactions with the disulfide bonds and the D-Trp side chains. One end of the PEG wraps around D-Lys8. D-Lys8 adopts a different conformation compared to Lys8 in the other 5 copies of Ro1 because of the favorable interactions with the PEG, and, because E-Tyr3 packs against D-Lys8, constraining it an orientation different than Lys8 of the other copies of Ro1. D-D-Trp7 adjusts to the unique orientation of D-Lys8 in order to maintain a stacking arrangement with D-Lys8, despite there being no significant constraints on the side chain orientation of D-D-Trp7, suggesting that the D-Trp7-Lys8 side-by-side stacking arrangement at the apex of the cyclic loop is a strongly favored interaction inherent to Ro1. Figures generated with the program Pymol (version 1.8.4.1).

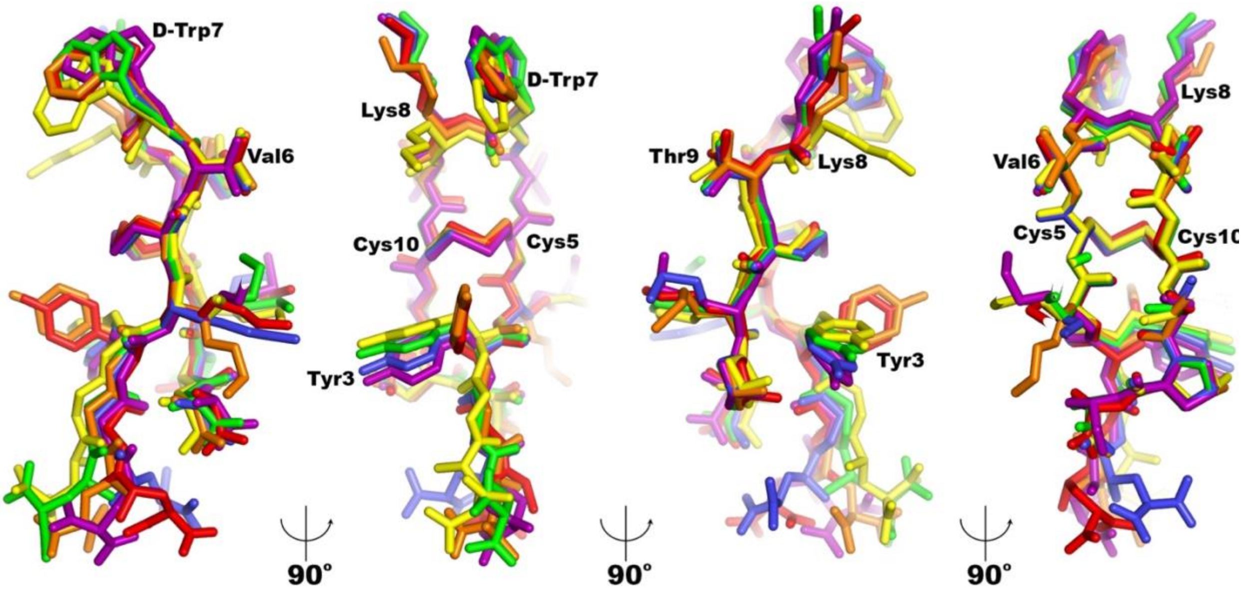


**Fig. S6.** Four views of the overlap of all copies of Consomatin Ro1 on copy **E** (colored orange). Alignment of the six copies of the molecule was performed with the program LSQKAB (*75*). Some of the residues are labeled in black. The side chain orientations of D-D-Trp7 and D-Lys8 differ from the other five copies yet the side-by-side stacking arrangement of D-Trp7 and Lys8 is a distinctive feature in all 6 copies of Ro1 that form the asymmetric unit of the crystal.

**Table S1.** Timeline of predation events for the 3 movie files provided for *Conus neocostatus* (Movies S3-S5).

| Time (h:min:sec) | Events |
|---|---|
| ***Movie S3*** | *Conus neocostatus* catches a fang blenny, single strike |
| 00:00:40 | Snail envenomates fish |
| 00:15:00 | Fish loses motor control; abnormal swimming behavior |
| 00:16:30 | Fish comes to rest on substrate |
| 01:15:00 | Snail starts tracking down fish |
| 01:22:00 | Fish appears to have died |
| 01:23:00 | Snail eats dead fish |
| ***Movie S4*** | *Conus neocostatus* envenomates a goby, night camera, two strikes |
| 00:00:00 | *Conus neocostatus* approaches fish from distance and strikes the first time |
| 00:16:00 | Goby starts climbing the wall and shows beginning of loss of balance |
| 00:35:00 | Snail becomes active, presumably stalking the fish |
| 01:02:00 | Fish appears to have recovered and is correctly balanced |
| 03:17:00 | Snail strikes a second time |
| 03:18:00 | Snail starts ingesting the fish shortly after the second strike |
| 03:19:00 | Final twitch from live fish while being ingested |
| ***Movie S5*** | *Conus neocostatus* catches a blenny with two strikes |
| 00:00:00 | Proboscis emerges |
| 00:02:02 | Snail envenomates fish (first sting) |
| 00:13:50 | Proboscis emerges again |
| 00:23:02 | Snail envenomates fish (second sting with brief tether) |
| 00:23:40 | Fish begins to climb wall |
| 00:24:20 | Fish falls and remains immobile |
| 01:06:00 | Fish appears motionless, possibly dead; snail engulfs the fish |

**Table S2.** Summarized activity of *C. rolani* venom fractions and synthetic Consomatin Ro1 in mice [intracranial (IC)]

| Age (days) | Weight (g) | n | Fraction/ Dose (nmol) | Observed Behavior (time = min post injection) |
|---|---|---|---|---|
| 15 | 7.33 | 1 | Control | normal: moving, responsive to prodding \|\| grooming, walking, rearing |
| | 7.84 | 1 | L516 16 | loss of balance at 3 min;<br>unresponsive to prodding at 17 min and lasts for at least 3.5 h<br>*recovered the next day, checked at ~18h |
| 16 | 8.38 ± 1.79 | 3 | Control | normal: moving, responsive to prodding \|\| grooming, walking, rearing |
| | 8.00 ± 0.42 | 2 | L516 16-12 (Consomatin Ro1 native) | initially normal: moving, responsive to prodding<br>hypoactive: less responsive to prodding 37 min - 1 h 5 min<br>heavy body, moves in place only, leans left<br>unresponsive: no movement when prodded except the tail 1h 8 min - 3 h<br>hypoactive: less responsive to prodding 3 h 18 min - 4 h 43 min<br>stretches only when prodded, moves in place, reacts sluggishly<br>mouse 1: checked the next day, recovered<br>mouse 2: observation ended at ~5h |
| Synthetic Consomatin Ro1 | | | | |
| 12 | 8.20 ± 0.11 | 2 | 0 | normal: moving, responsive to prodding \|\| grooming, walking, rearing |
| | 7.57± 0.14 | 2 | 10 | less responsive to non-responsive to prodding from 1 h 15 min and lasted for 3 h 5 min |
| | 8.09 | 1 | 25.75 | less responsive to non-responsive to prodding from 31 min and lasted for 4 h 15 min |

**Table S2 (Continued).** Summarized activity of *C. rolani* venom fractions and synthetic Consomatin Ro1 in mice [intracranial (IC)]

| Synthetic Consomatin Ro1 | | | | |
|---|---|---|---|---|
| 16 | 7.33 ± 0.38 | 6 | 0 | normal: moving, responsive to prodding \|\| grooming, walking, rearing |
| | 7.72 ± 0.21 | 2 | 2.5 | (only one out of 2 mice tested showed activity)<br>less responsive to prodding from 2 h and lasted for 2 h 6 min<br>recovered at 4 h |
| | 7.25 ± 0.06 | 2 | 5 | less responsive to non-responsive to prodding from 3 h 11 min and lasted for 1 h 11 min<br>*left overnight<br>*recovered the next day, checked at ~18-19 h |
| | 8.14 ± 0.18 | 2 | 10 | less responsive to non-responsive to prodding from 2 h 4 min and lasted for 1.5 h |
| | 7.58 ± 0.81 | 2 | 20 | less responsive to non-responsive to prodding from 1 h 4 min and lasted for 3 h 37 min<br>*left overnight<br>*recovered the next day, checked at ~17 h |
| | 7.25 | 1 | 30 | less responsive to prodding from 1 h 24 min and lasted for 2 h 43 min |
| 17 | 9.10 ± 0.70 | 3 | 0 | normal: moving, responsive to prodding \|\| grooming, walking, rearing |
| | 7.39 | 1 | 1 | normal: grooming, moving, responsive to prodding |
| | 7.54 ± 0.16 | 2 | 2.5 | less responsive to prodding from 2 h 2 min and lasted for 2 h |
| | 8.97 ± 0.73 | 2 | 10 | less responsive to prodding from 1 h 25 min and lasted for 1 h 39 min |
| | 9.13 ± 0.18 | 2 | 12.38 | less responsive to non-responsive to prodding from 1h 45 min and lasted for 2 h 19 min |
| | 9.17 | 1 | 25.75 | less responsive to non-responsive to prodding from 2 h 7 min and lasted for 1 h 53 min |
| 19 | 10.15 ± 0.84 | 5 | 0 | normal: moving, responsive to prodding \|\| grooming, walking, rearing |
| | 9.45 ± 0.57 | 2 | 5 | less responsive to non-responsive to prodding from 2 h 12 min and lasted for 2 h 30 min |
| | 10.27 | 1 | 30 | less responsive to non-responsive to prodding from 2 h<br>*assay cut short at 2h 30 min |
| 22 | 14.53 ± 0.57 | 2 | 0 | normal: moving, responsive to prodding \|\| grooming, walking, rearing |
| | 13.61 | 1 | 20 | less responsive to prodding from 4 h and lasted for 3 h |

**Table S3.** Consomatin Ro1 crystallographic data and refinement statistics

| **Data** | | |
|---|---|---|
| Crystal | WT | $K_2PtCl_4$-soak |
| Source/Wavelength/Date of data collection | SSRL 9-2 / 1.7711/ 05/12/2019 | SSRL 14-1 / 1.0716/ 07/16/2019 |
| Space Group (unit cell dimensions, Å) | $P6_522$ (50.28, 50.28, 135.01) | $P6_522$ (50.32, 50.32, 134.60) |
| Resolution (Å) | 36.59 – 1.95 | 36.58 – 2.40 |
| Resolution (Å) (high-resolution shell) | (2.00 – 1.95) | (2.49 – 2.40) |
| # Reflections measured | 1,125,051 | 615,190 |
| # Unique reflections | 8,016 | 4,434 |
| Redundancy (high-resolution shell) | 140 (103) | 139 (144) |
| Completeness (%)(high-resolution shell) | 99.9 (98.2) | 100.0 (99.7) |
| Anomalous redundancy (high-resolution shell) | | 82 (80) |
| Anomalous completeness (%)(high-resolution shell) | | 99.9 (99.6) |
| <I/σI> (high-resolution shell) | 24.3 (1.4) | 24.8 (3.7) |
| <CC1/2> | 1.000 (0.700) | 0.999 (0.906) |
| Rpim (high-resolution shell) | 0.019 (0.744) | 0.023 (0.389) |
| Mosaicity (°) | 0.11 | 0.13 |
| **Refinement** | | |
| Program | Phenix.refine | |
| Resolution (Å) | 36.62 – 1.95 | |
| Resolution (Å) – (high-resolution shell) | (2.01 – 1.95) | |
| # Reflections used for refinement | 6,796 | |
| # Reflections in Rfree set | 1,167 | |
| Rcryst | 0.239 (0.304) | |
| Rfree | 0.283 (0.379) | |
| RMSD: bonds (Å) / angles (°) | 0.011 / 2.068 | |
| <B> ($Å^2$): all atoms / # atoms | 61.0 / 723 | |
| <B> ($Å^2$): water molecules / #water | 54.2 / 34 | |
| <B> ($Å^2$): PEG molecule / #non-hydrogen atoms ($C_{34}O_{17}H_{69}$) | 62.7 / 51 | |
| ϕ/ψ most favored (%) / additionally allowed (%) | 100.0/0.0 | |

**Table S4.** Molar $EC_{50}$ values for human SS-14, Consomatin Ro1, Consomatin G1 at the human $SST_{1-5}$ using the PRESTO-Tango

| | Human SS-14 | Consomatin Ro1 | Consomatin G1 |
|---|---|---|---|
| **$SST_1$** | | | |
| $EC_{50}$ | 3.72E-09 | 2.88E-06 | |
| $pEC_{50}$ | 8.43 | 5.54 | |
| +/-CI95 | 0.39 | 0.77 | |
| +/- SEM | 0.16 | 0.18 | |
| n | 7 | 3 | |
| **$SST_2$** | | | |
| $EC_{50}$ | 1.26E-08 | | 2.58E-09 |
| $pEC_{50}$ | 7.90 | | 8.59 |
| +/-CI95 | 0.22 | | 0.58 |
| +/- SEM | 0.09 | | 0.13 |
| n | 7 | | 3 |
| **$SST_3$** | | | |
| $EC_{50}$ | 4.58E-08 | | |
| $pEC_{50}$ | 7.34 | | |
| +/-CI95 | 0.29 | | |
| +/- SEM | 0.12 | | |
| n | 7 | | |
| **$SST_4$** | | | |
| $EC_{50}$ | 4.54E-09 | 5.05E-06 | |
| $pEC_{50}$ | 8.343 | 5.30 | |
| +/-CI95 | 0.522 | 0.28 | |
| +/- SEM | 0.213 | 0.06 | |
| n | 7 | 3 | |
| **$SST_5$** | | | |
| $EC_{50}$ | 3.00E-08 | | |
| $pEC_{50}$ | 7.52 | | |
| +/-CI95 | 0.36 | | |
| +/- SEM | 0.15 | | |
| n | 7 | | |

**Table S5.** Venom gland transcriptome datasets interrogated in this study for the bioinformatic identification of SS-like sequences

| *Conus* species | Subgenera | No of assembled contigs | Accession numbers |
|---|---|---|---|
| *Conus ermineus* | *Chelyconus* | 7,543 | SRA: SRR6983169 |
| *Conus marmoreus* | *Conus* | 18,596 | SRA: SRX5015020, SRX1323884 |
| *Conus gloriamaris* | *Cylinder* | 15,437, 14,178 | SRA: SRX2779517, SRR5499408 |
| *Conus textile* | *Cylinder* | 19,161 | SRA: SRX5015023 |
| *Conus episcopatus* | *Darioconus* | 5,767 | DDBJ: SAMD00029746 |
| *Conus betulinus* | *Dendroconus* | 32,400 | SRA: SRR2124881 |
| *Conus geographus* | *Gastridium* | 9,191 | SRA: SRX151242, SRX151241, SRX151240, SRX151239 |
| *Conus sponsalis* | *Harmoniconus* | 3,700 | SRA: SRX1323890 |
| *Conus trochulus* | *Kalloconus* | 18,552 | SRA: SRR11807506 |
| *Conus antoniomonteiroi* | *Lautoconus/Africonus* | 26,034 | SRA: SRR11807494 |
| *Conus boavistensis* | *Lautoconus/Africonus* | 6,882 | SRA: SRR11807497 |
| *Conus cuneolus* | *Lautoconus/Africonus* | 4,220 | SRA: SRR11807496 |
| *Conus galeao* | *Lautoconus/Africonus* | 8,962 | SRA: SRR11807500 |
| *Conus grahami* | *Lautoconus/Africonus* | 10,682 | SRA: SRR11807507. |
| *Conus infinitus* | *Lautoconus/Africonus* | 12,104 | SRA: SRR11807493 |
| *Conus maioensis* | *Lautoconus/Africonus* | 45,991 | SRA: SRR11807501, SRR11807499 |
| *Conus miruchae* | *Lautoconus/Africonus* | 12,992 | SRA: SRR11807495 |
| *Conus raulsilvai* | *Lautoconus/Africonus* | 18,028 | SRA: SRR11807492 |
| *Conus verdensis* | *Lautoconus/Africonus* | 14,942 | SRA: SRR11807498 |
| *Conus guanche* | *Lautoconus/Varioconus* | 17,586 | SRA: SRR11807502 |
| *Conus mercator* | *Lautoconus/Varioconus* | 27,651 | SRA: SRR11807503, SRR11807505 |
| *Conus reticulatus* | *Lautoconus/Varioconus* | 11,242 | SRA: SRR11807504 |
| *Conus lividus* | *Lividoconus* | 3,951 | SRA: SRX1323888 |
| *Conus quercinus* | *Lividoconus* | 13,002 | CNGB: CNS0048932 |
| *Conus magus* | *Pionoconus* | 45,382 | SRA: SRX5015024, SRR9831255 |
| *Conus striatus* | *Pionoconus* | 29,445 | SRA: SRX5015022 |
| *Conus arenatus* | *Puncticulis* | 3,581 | SRA: SRX1323893 |
| *Conus caracteristicus* | *Puncticulis* | 20,037 | CNGB: CNS0048931 |
| *Conus rattus* | *Rhizoconus* | 3,982 | SRA: SRX1323889 |
| *Conus bayani* | *Splinoconus* | 43,997 | SRA: SRR13781584 |
| *Conus imperialis* | *Stephanoconus* | 2,690 | SRA: SRX1323891 |
| *Conus generalis* | *Strategoconus* | 25,594 | CNGB: CNS0048933 |
| *Conus varius* | *Strategoconus* | 5,606 | SRA: SRX1323892 |
| *Conus terebra* | *Virgiconus* | 11,520 | SRA: SRX5015025 |
| *Conus virgo* | *Virgiconus* | 25,178 | SRA: SRX1323883, SRX5015021 |
| *Conus coronatus* | *Virroconus* | 2,715 | SRA: SRX1323894 |
| *Conus ebraeus* | *Virroconus* | 3,856 | SRA: SRX1323887 |

**Movie S1**: The taser-and-tether hunting strategy. *Conus bullatus* (type species of the *Textilia* clade) catching a blenny.

**Movie S2**: The net hunting strategy. *Conus geographus* (type species of the *Gastridium* clade) chasing after and catching a group of fish.

**Movie S3**: The ambush-and-assess hunter *Conus neocostatus* envenomates a fang blenny, *Meiacanthus grammistes*. Fish dies 1:15 h after the first sting.

**Movie S4**: The ambush-and-assess hunter *Conus neocostatus* envenomates a fish (genus Amblyeleotris) with two strikes, caught on night camera.

**Movie S5**: The ambush-and-assess hunter *Conus neocostatus* catches a blenny with two strikes.

**Movie S6:** *Conus bullatus* being attacked by an aggressive fish (type species of the *Textilia* clade).
